# Sex-specific differences in brain activity dynamics of youth with a family history of substance use disorder

**DOI:** 10.1101/2024.09.03.610959

**Authors:** Louisa Schilling, S Parker Singleton, Ceren Tozlu, Marie Hédo, Qingyu Zhao, Kilian M Pohl, Keith Jamison, Amy Kuceyeski

**Affiliations:** Department of Radiology, Weill Cornell Medicine, New York, NY, USA; Department of Psychiatry & Behavioral Sciences, Stanford University School of Medicine, Stanford, California, USA

**Keywords:** substance use disorder, family history, adolescence, sex differences, neuroimaging, network control theory, brain activity dynamics

## Abstract

An individual’s risk of substance use disorder (SUD) is shaped by a complex interplay of potent biosocial factors. Current neurodevelopmental models posit vulnerability to SUD in youth is due to an overreactive reward system and reduced inhibitory control. Having a family history of SUD is a particularly strong risk factor, yet few studies have explored its impact on brain function and structure prior to substance exposure. Herein, we utilized a network control theory approach to quantify sex-specific differences in brain activity dynamics in youth with and without a family history of SUD, drawn from a large cohort of substance-naïve youth from the Adolescent Brain Cognitive Development Study. We summarize brain dynamics by calculating transition energy, which probes the ease with which a whole brain, region or network drives the brain towards a specific spatial pattern of activation (i.e., brain state). Our findings reveal that a family history of SUD is associated with alterations in the brain’s dynamics wherein: i) independent of sex, certain regions’ transition energies are higher in those with a family history of SUD and ii) there exist sex-specific differences in SUD family history groups at multiple levels of transition energy (global, network, and regional). Family history-by-sex effects reveal that energetic demand is increased in females with a family history of SUD and decreased in males with a family history of SUD, compared to their same-sex counterparts with no SUD family history. Specifically, we localize these effects to higher energetic demands of the default mode network in females with a family history of SUD and lower energetic demands of attention networks in males with a family history of SUD. These results suggest a family history of SUD may increase reward saliency in males and decrease efficiency of top-down inhibitory control in females. This work could be used to inform personalized intervention strategies that may target differing cognitive mechanisms that predispose individuals to the development of SUD.

## 1 Introduction

Substance use disorder (SUD) frequently leads to escalating consequences including familial and financial instability, poor health outcomes, and – far too often – death (1). However, only a subset of those regularly exposed to substances of abuse go on to develop SUD (2), highlighting the critical need to identify predisposing factors. Consistent with the known genetic component of SUD (2; 3), youth with a family history of SUD (FH+) are at higher risk of SUD than youth without a family history of SUD (FH-) (4). This may be due to an exacerbated developmental profile of adolescence, characterized by poor inhibitory control and a highly sensitive reward system (4; 5; 6; 7). Additionally, parental history of SUD is itself considered an adverse childhood experience and thus poses both environmental and genetic risks (8).

Recent work suggests that structural and functional signatures of SUD partly reflect genetically and environmentally-conferred predisposition for SUD, rather than sole consequences of chronic substance use (9; 10; 11). Extensive work has revealed individuals with SUD have altered dopaminergic signaling (12), white matter integrity (13; 14), functional activity (14; 15; 16), gray matter volume (14; 17), and network dynamics (18; 19). These findings are largely localized to brain regions involved in reward and inhibitory control such as the basal ganglia and prefrontal cortex (20; 21). Lagging slightly behind is a growing body of parallel findings in individuals at high risk of SUD, including altered dopaminergic signaling (22), white matter integrity (23; 24; 25), functional activity (6; 26), gray matter volume (27), and network dynamics (19; 28; 29) also primarily in regions responsible for behavioral inhibition and reward processing (5; 6). However, this remains an emerging area of research with few well-powered studies in youth with no previous substance exposure. As substance use impacts the brain’s structure and function, studying FH+ youth *prior* to substance exposure is crucial for disentangling the driving forces from the consequences of problematic substance use.

Sex assigned at birth (hereafter, “sex”) modulates SUD at every phase of the disorder (e.g., onset, manifestation, withdrawal, craving, relapse), likely due to a confluence of biological and sociocultural correlates of sex and gender (30; 31; 32; 33). Multiple lines of evidence converge on a model in which females are more susceptible to negative reinforcers (e.g., withdrawal, stress) and males are more susceptible to positive reinforcers (e.g., pleasurable effects of drugs) of SUD (32; 34; 35; 36; 37; 38). While men are more likely to use illicit drugs and develop SUD, women escalate drug use more quickly (i.e., “telescoping” effect) and report more withdrawal and craving (32). Moreover, females are more likely to have an internalizing pathway and males more likely to have an externalizing pathway to SUD (35; 36; 37; 38). Further, sex differences in the effect of family history on adolescents have been observed in behavioral (39; 40; 41; 42) and neurobiological phenotypes (43). Yet, few neuroimaging studies have considered whether sex modifies the effect of family history of SUD.

Herein, we aim to characterize sex-specific differences in the brain’s functional activation dynamics in substance-naïve youth from the Adolescent Brain Cognitive Development (ABCD) Study(44). Resting-state functional MRI (rsfMRI) collected in these adolescents allows tracking of brain activity as it alternates between several commonly recurring states (45; 46). Network control theory (NCT) can be applied to model how the brain dynamically moves through these states (47; 48). NCT considers the brain to be a network of regions interconnected via white matter tracts (i.e. structural connectivity) and models brain activity through a linear time-invariant system that assumes functional activation flows via a network diffusion process. The sum of activational inputs applied during network diffusion to steer or *control* a trajectory from an initial state to a desired target state represent *transition energy* (TE)(48). In the current context, TE inputs can be thought of as the internal demands required to drive the brain towards a given brain state (49; 50), and does not necessarily reflect metabolic demands, though recent work has found TE to be related to metabolic activity (51; 52). NCT-based analyses have found TE to be i) correlated with age (53), ii) increased in heavy alcohol use in young adults (18), iii) related to neurotransmitter receptor function (50), iv) altered in psychopathology (54), and v) change with pharmacological intervention (55). The relationship between TE and a familial predisposition to SUD in youth remains unknown. We hypothesize that FH+ youth compared to FH-youth will exhibit i) increased global TE, ii) network and regional energetic differences in areas involved in cognitive control and reward sensitivity, and, finally, iii) that these differences will be sex-specific.

## 2 Results

### 2.1 Sample Characteristics

To investigate sex-specific differences in the energetic landscapes of youth with and without a family history of SUD, we use diffusion and resting state functional MRI (rsfMRI) data from the ABCD Study’s baseline assessment of a large sample of substance-naïve youth (N= 1894 individuals, ages 10.02 ± 0.63, 54% female) (Table 1). See Section 5.1.1 for exclusion criteria. We classify individuals as FH+ if they have at least one parent or two grandparents with a history of SUD and FH-if no parents or grandparents had a history (56; 57; 58). Individuals with just one grandparent with a history of SUD are classified as FH+/- and are included only in analyses of continuous associations (i.e., family history density).

**Table 1.**
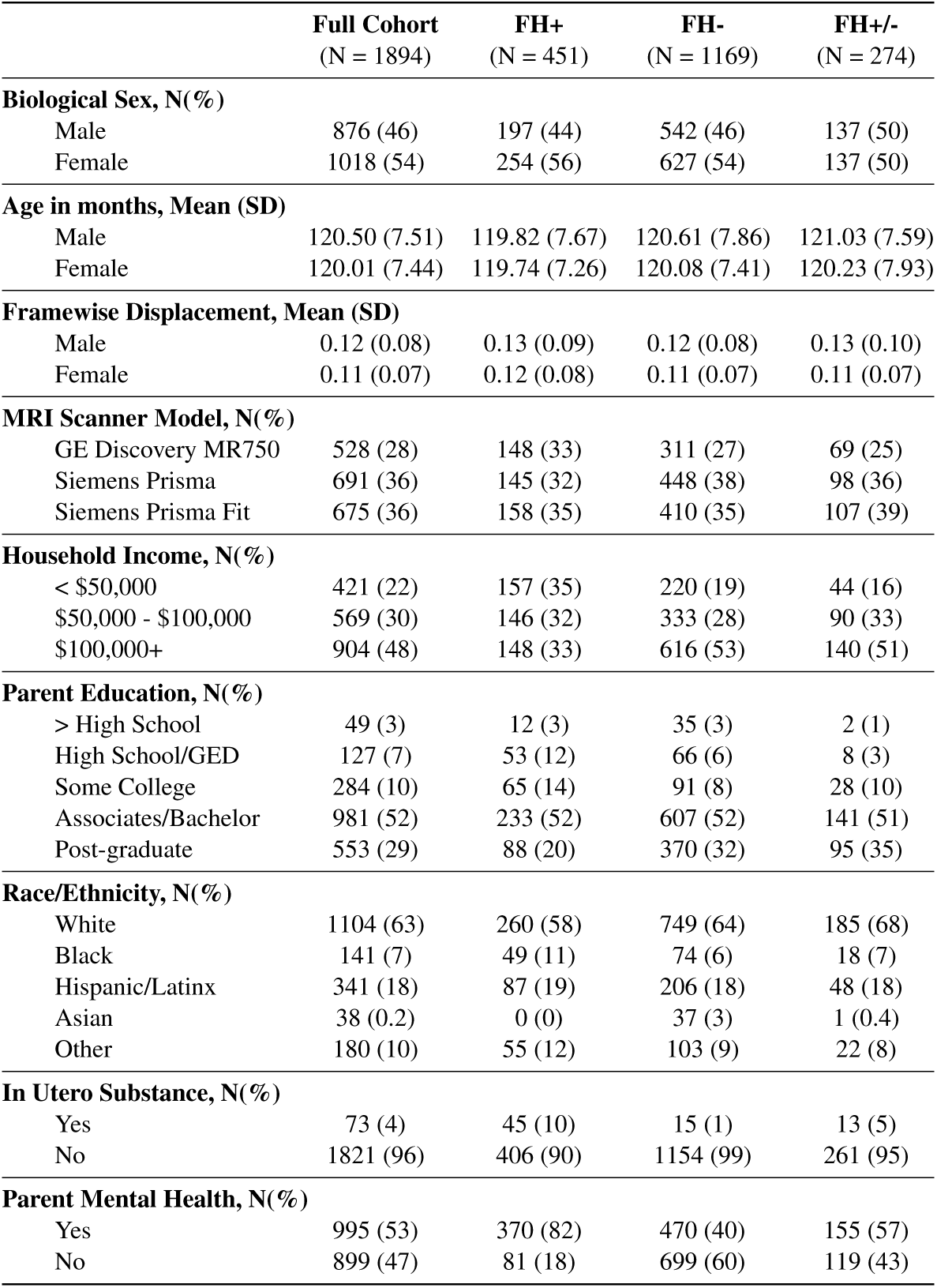
Demographic and characteristic data for the full cohort and subgroups (FH+, FH-, and FH+/-).

### 2.2 Network Control Theory Analysis

Following previous work (47; 55), we apply *k*-means clustering to regional rsfMRI time series data (86-region atlas) to identify *k* recurring patterns of brain activity, termed “brain states” (Figure 1, a-c). For each subject, we assign each individual frame to a brain state and calculate subject-specific brain state centroids (Figure 1, d-e). We then apply NCT using a group-average structural connectome (from a subset of individuals in this dataset) to calculate the global, network, and region-level TE required to complete brain state transitions (Figure 1, f-g). At the global, network and region-levels, we calculate pairwise and mean TE (Figure 1g). Pairwise TE represents the energy a given region, network or whole brain requires to complete transitions between each pair of brain states, whereas mean TE represents the overall energetic demands of a region, network or whole brain to move through its state space (i.e., across all pairs of states). Initial TE calculations result in “pairwise regional TE” values, i.e., 86 (number of regions) *k*x*k* matrices. For each matrix, we sum pairwise regional TE of all regions belonging to each of the seven Yeo networks (59) plus subcortical and cerebellar networks to yield “pairwise network TE” values, i.e., 9 (number of networks) *k*x*k* matrices. We derive a single “pairwise global TE” *k*x*k* matrix by summing across all 86 regions pairwise TEs for each transition *k*x*k* matrix. We also average all entries in pairwise TE matrices to derive mean TE values for each region, network and globally, resulting in: 86 mean regional TE values, 9 mean network TEs, and a single value for mean global TE. See section 5.3.2 for more details.

**Figure 1.**
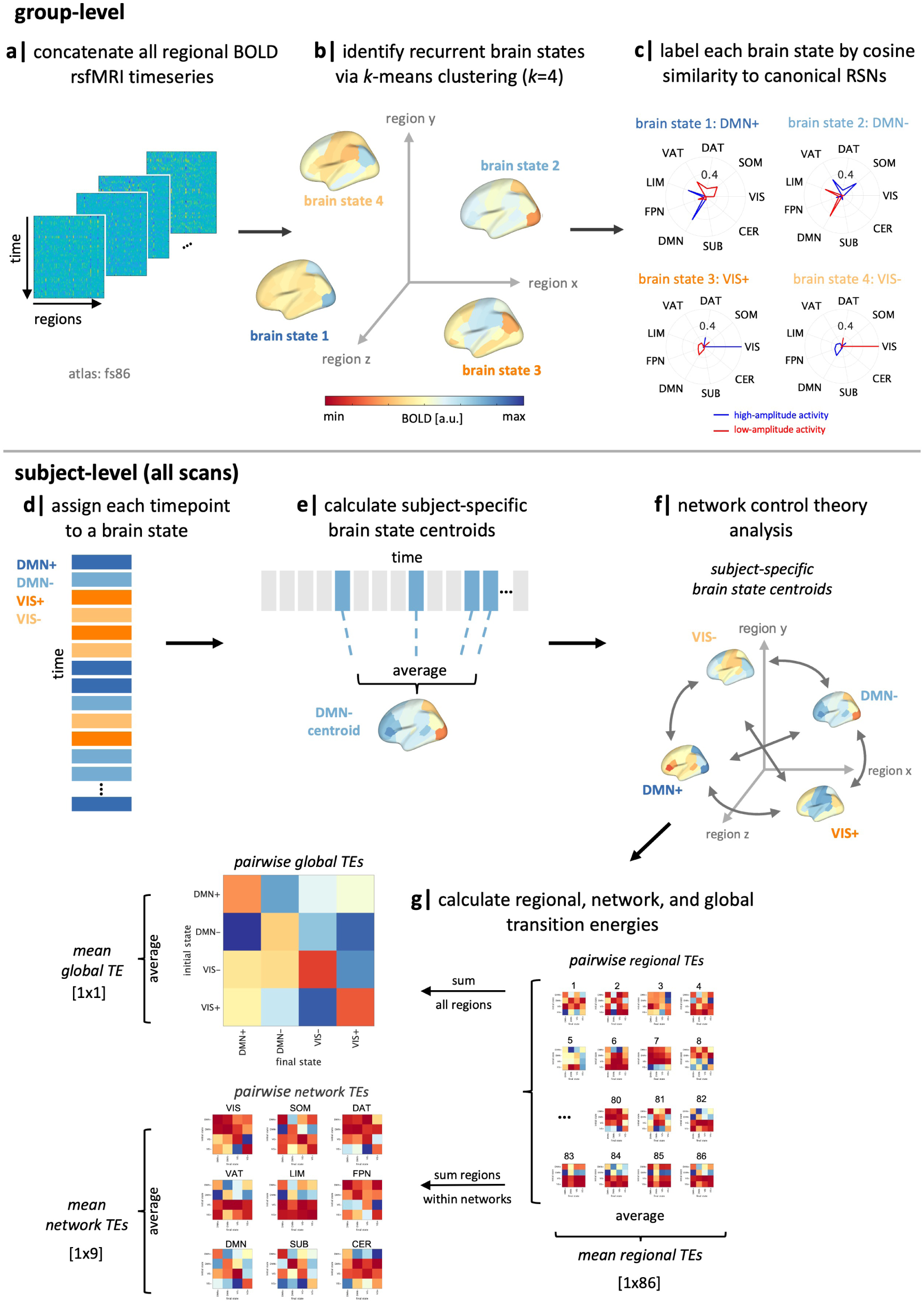
Workflow. (a-b) We applied *k*-means clustering to regional rsfMRI time series of all subjects to identify four recurring “brain states”. (c) We calculated the cosine similarity of high and low amplitude activity with the Yeo 7-networks (59) plus subcortical and cerebellar networks and named each state by taking the maximum of those values. (d-e) For each subject, individual time frames in the fMRI scan were assigned to a brain state, and subject-specific brain state centroids were calculated. (f) Network control theory was then implemented using a group-average structural connectome to calculate the transition energy (TE) required for transitioning between pairs of subject-specific brain states. (g) Pairwise and mean TE values were computed at global, network, and regional levels for every individual in the dataset. SUB = subcortical structures, CER = cerebellar structures, VIS = visual network, SOM = somatomotor network, DAT = dorsal attention network, VAT = ventral attention network, LIM = limbic network, FPN = frontoparietal network, DMN = default mode network, RSN = resting-state network, TE = transition energy.

It is important to distinguish between the four identified brain states and the nine networks for which we calculate TE values, though they are related to a common network parcellation of the 7 Yeo networks plus subcortical and cerebellar networks. Brain states refer to the four clusters identified by *k*-means clustering, which are each assigned to one of the nine networks based on the network whose isolated high or low amplitude activity best explain each brain state. We refer to them as “DMN+”, “DMN-”, “VIS+”, and “VIS-”, using the network name and respective indicators representing activity above (+) or below (-) regional means in the the DMN and VIS networks. These names are purely descriptive and did not influence analyses. Network TE values, on the other hand, refer to the TE values calculated by summing across regional TE values for regions assigned to each network. We reference these using the nine network names alone, e.g. “DAT”, “VIS”, “DMN”, etc.

We run a series of analysis of covariances (ANCOVAs) to investigate the effect of family history of SUD and its interaction with sex on mean and pairwise TE at the global, network, and regional levels. We include sex, age, family history of SUD (FH+ vs FH-), race/ethnicity, household income, parental education, parental mental health issues, in utero exposure to substances, MRI scanner model, in-scanner motion (i.e., mean framewise displacement), and two interaction terms: sex and family history of SUD, and income and family history of SUD as independent variables in the model. To determine the direction of effect between groups, we performed post hoc *t* tests on TE values found to be significant in ANCOVA models.

### 2.3 Brain State Identification

We found an optimal *k*=4 brain states, as determined using the additional explained variance <1% cut-off (Figure S1). The identified brain states consist of two pairs of anti-correlated activity patterns (i.e. meta-states), the first dominated by high and low-amplitude activity in the default mode network (DMN+/-), and the second by high and and low-amplitude activity in the visual network (VIS+/-) (Figure 2), aligning with previous work (18; 47). For replication of our results using *k*=5, see S8.

**Figure 2.**
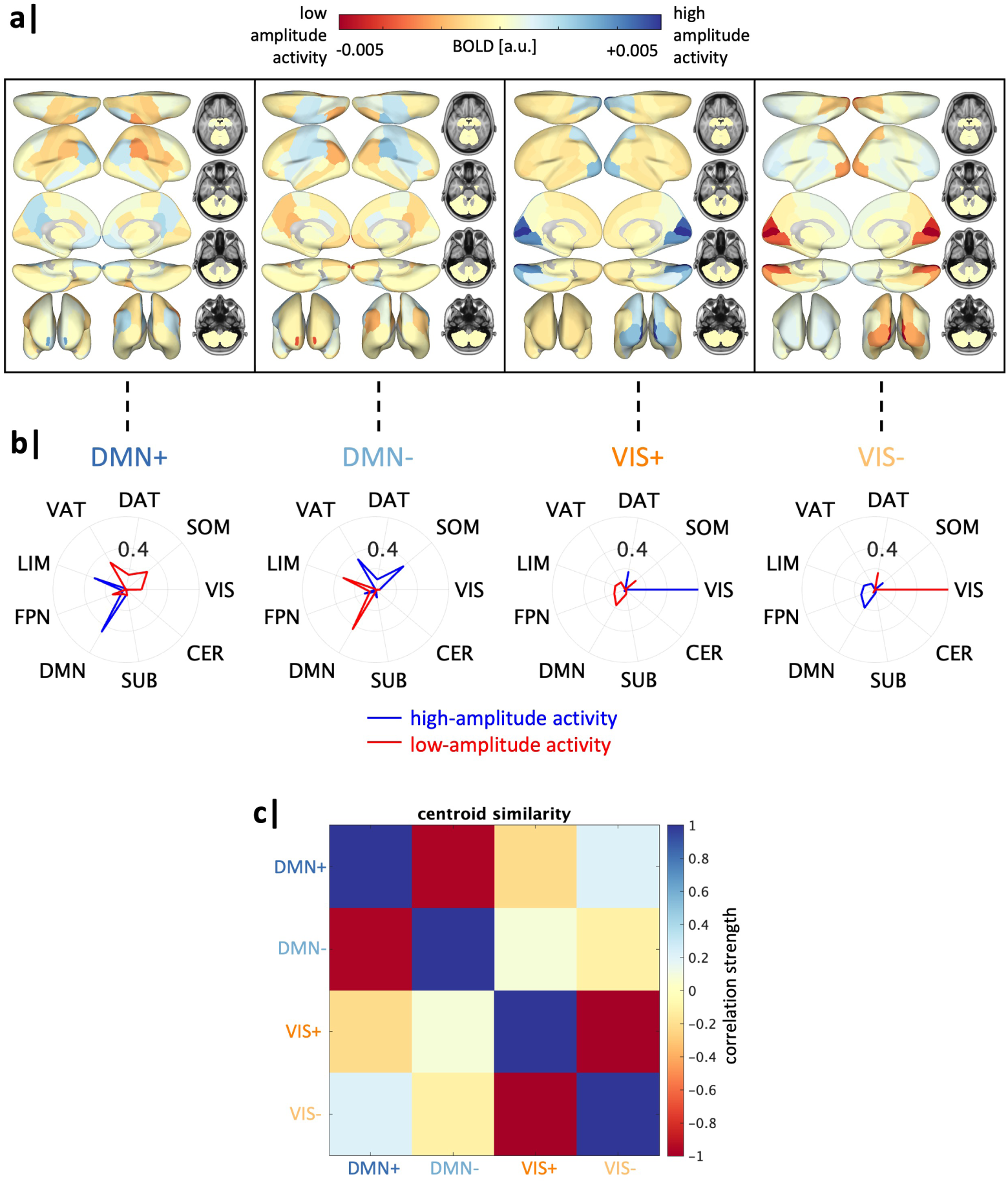
Global transition energy differences in family history of SUD vary by sex. Four recurrent states of brain activity (brain states) identified via *k*-means clustering across all subjects. Group-average centroids are used for all illustrations. (a) Mean BOLD activation of each brain state plotted on the brain’s surface. a.u. = arbitrary units. (b) Cosine similarity with canonical resting-state networks (59) was calculated for the positive (high-amplitude) and negative (low-amplitude) components of each brain state’s group-average centroid separately. Each brain state is assigned the label of the network with the maximal cosine similarity value, with a sign indicating the maximal similarity was determined using high amplitude activity (+) or low amplitude activity (-). (c) Pearson correlation values between each pair of brain states. a.u. = arbitrary units, SUB = subcortical structures, CER = cerebellar structures, VIS = visual network, SOM = somatomotor network, DAT = dorsal attention network, VAT = ventral attention network, LIM = limbic network, FPN = frontoparietal network, DMN = default mode network.

### 2.4 Global Transition Energy

We first sought to determine whether a family history of SUD and its interaction with sex have an observable global effect on the brain’s overall energetic landscape. In an ANCOVA model for mean global TE, including the covariates described above, the effect of family history of SUD was not significant; however, the interaction of sex and family history of SUD was significant (Table 2). The lack of an effect of family history of SUD on global TE is partially due to its interaction with sex, wherein an opposite effect of family history is observed in females and males. To determine the direction of this sex-specific effect, we performed within-sex unpaired *t* tests comparing FH+ and FH-individuals. FH+ females exhibit a significantly higher mean global TE compared to FH-females (*t* = 2.14, *p* = 0.03, *p*FDR = 0.04), whereas FH+ males exhibit a non-significant trend toward lower mean global TE compared to FH-males (*t* = -1.03, *p* = 0.30) (Figure 3a). Further, when including FH+/- individuals (i.e., subjects with 1 grandparent with SUD), mean global TE and family history density of SUD were found to have a weak positive trend in females (rho = 0.07, *p* = 0.07), while males were not correlated (rho = -0.03, *p*= 0.41) (Figure 3b).

**Figure 3.**
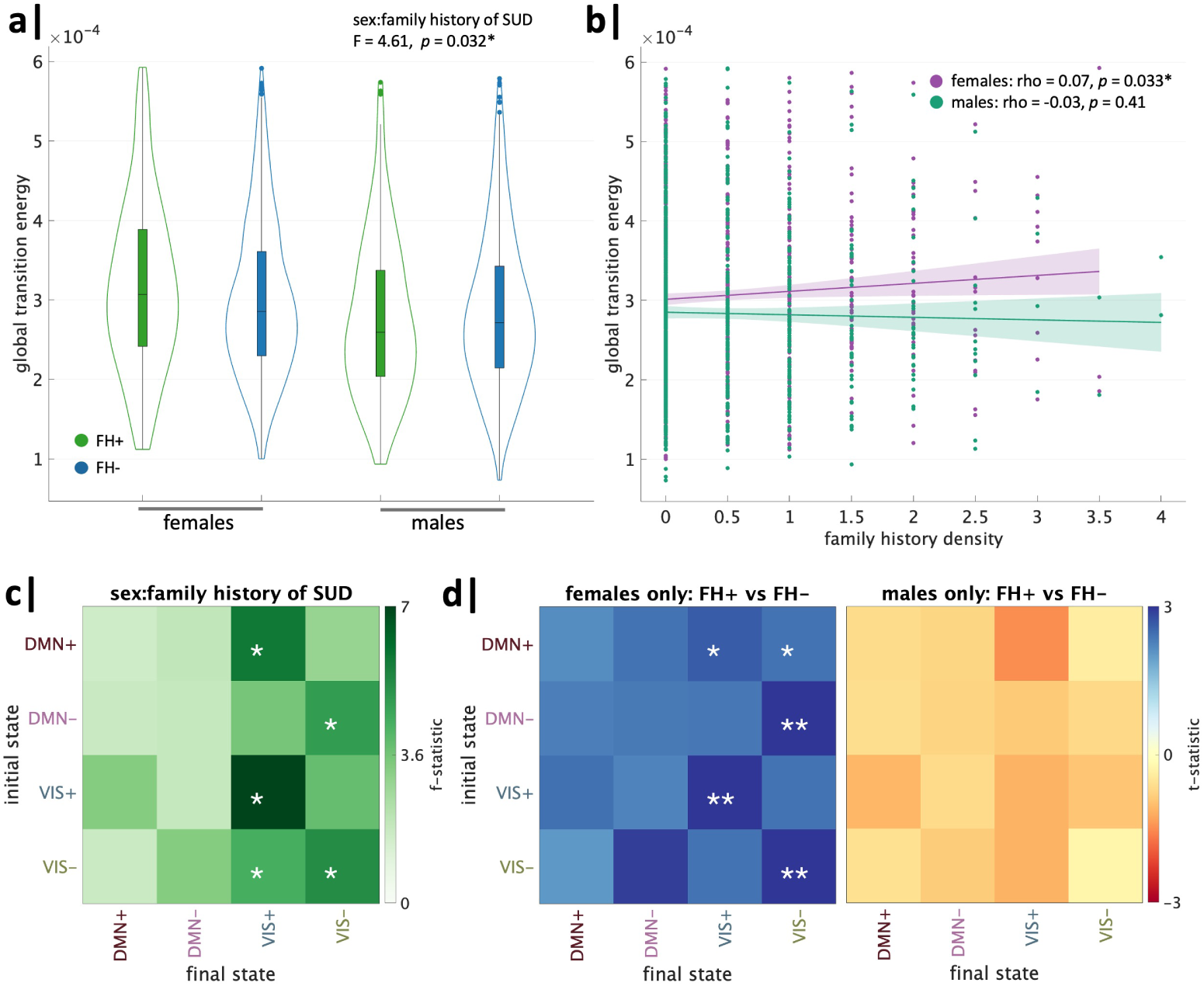
Global transition energy differences in family history of SUD vary by sex. Analysis of global transition energy revealed a significant effect of family history of SUD by sex, wherein (a) FH+ females > FH-females and FH+ males < FH-males, (b) family history density had a trend toward a weak positive correlation with global TE in females and no correlation in males. (c) Whole brain TE between pairs of states revealed the interaction between family history of SUD and sex was driven by transitions to/from visual network states. (d) Post-hoc t-tests revealed FH+ females had higher TE than FH-females across all transitions, particularly in persistence of visual network states, and FH+ males had lower TE than FH-males across all transitions (none significant). *significant before and **significant after correcting for the 16 (4×4) comparisons.

**Table 2.** Mean Global Transition Energy ANCOVA (FH+ vs. FH-)

| Variable | F-statistic | p |
| --- | --- | --- |
| Family History of SUD (FH+ vs FH-) | 0.17 | 0.70 |
| Age | 0.05 | 0.82 |
| Sex | 27.67 | 1.6e-07** |
| Family History of SUD : Sex | 4.61 | 0.032* |
| Household Income | 0.52 | 0.60 |
| Family History of SUD : Household Income | 4.34 | 0.013* |
| Parental Education | 0.13 | 0.87 |
| Race/Ethnicity | 0.42 | 0.80 |
| In Utero Substance Exposure | 0.019 | 0.89 |
| Parental History of Mental Health | 5.01 | 0.025* |
| Framewise Displacement | 0.65 | 0.42 |
| MRI Scanner Model | 21.93 | 4.03e-10** |

\* = p-value significant, \*\* = p-value significant after FDR correction

Notably, the effect size of sex on mean global TE was the largest out of all variables examined, with females exhibiting higher values compared to males (post hoc *t*= 4.7, *p* < 0.0001, *p*FDR < 0.0001). Children of parents with mental health issues exhibited higher global TE than those without parental mental health history (*t* = 3.08, *p* = 0.002, *p*FDR = 0.004). MRI scanner model and the interaction of family history of SUD and household income were also significant. See Supplemental Information for within-income category, within-scanner type, and single-site analyses (Figs. S12, S13, S14).

ANCOVA models for entries in the pairwise global TE matrix revealed the effect of the interaction of sex and family history of SUD was strongest in transitions to and, particularly, persistence within the VIS+/- brain states (Figure 3c). Post hoc unpaired t-tests revealed this interaction effect was primarily driven by increased pairwise global TE in FH+ females versus FH-females in transitions to the VIS+/- brain states (Figure 3d). All transitions had higher TE in FH+ females and lower TE in FH+ males compared to sex-matched FH-counterparts.

### 2.5 Network Transition Energy

We next investigated which networks may be driving the observed effects in global TE. For each network, we ran ANCOVAs for mean network TE using the same covariates described above. Mean network TE did not have a significant effect of family history of SUD alone in any network (Figure 4a), but significant interaction effects of sex and family history of SUD were found in the default mode (DMN), dorsal attention (DAT), and ventral attention (VAT) networks, though the VAT did not survive FDR correction (Figure 4b). ANCOVA results for all networks can be found in the Supplementary Information. To determine the direction of interaction effects in the DMN, DAT and VAT, we performed unpaired *t* tests to compare FH+ and FH-individuals within each sex. We found the family history-by-sex effect in the DMN was driven by higher mean network TE in FH+ females than FH-females (*t* = 2.44, *p* = 0.015, *p*FDR = 0.030) (Figure 4c). Further, family history density of SUD in females had a weak trending positive correlation with mean DMN TE (rho = 0.053, *r* = 0.090) (Figure 4f). On the other hand, family history-by-sex effects in the DAT and VAT were driven by decreased mean network TE in FH+ males compared to FH-males (DAT: *t* = -3.54, *p*FDR = 0.00025; VAT: *t* = -2.62, *p* = 0.0091, *p*FDR = 0.027) (Figure 4,d-e). There were also significant, though weak, negative correlations between mean network TE and FHD in males in DAT (males: rho = -0.103, *p* = 0.0022, *p*FDR = 0.011) and VAT (males: rho = -0.072, *p* =0.033, *p*FDR = 0.10), though the VAT correlation did not survive FDR correction (Figure 4,g-h).

**Figure 4.**
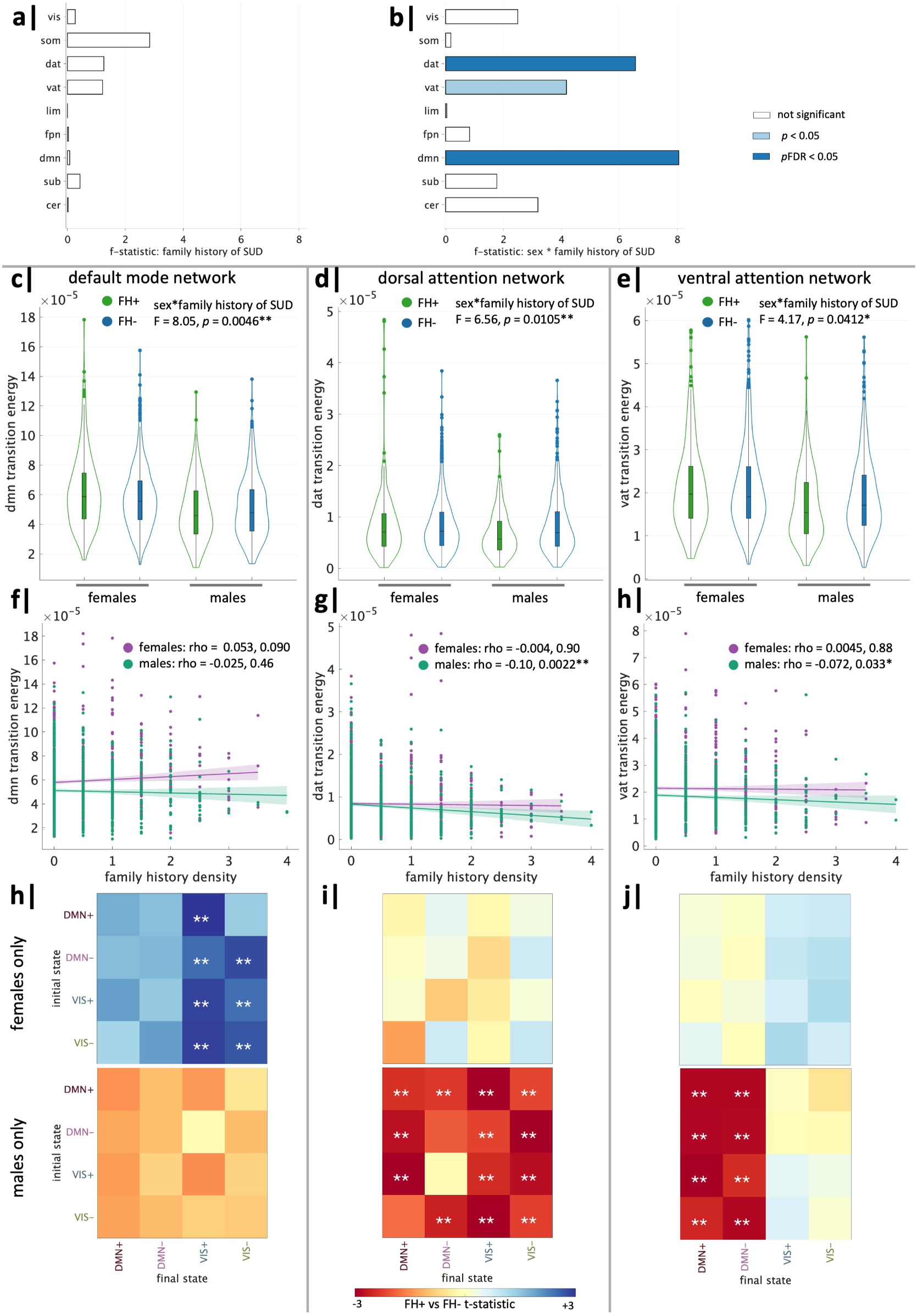
Network-level transition energy differences in family history of SUD vary by sex. ANCOVA analysis of mean network transition energy revealed (a) no significant effects of family history of SUD and (b) significant effects of family history of SUD by sex in the default mode (DMN), dorsal attention (DAT) and ventral attention (VAT) networks. DMN: (c) The interaction effect was driven by FH+ females > FH-females. (f) Family history density trended toward a weak positive correlation with DMN network TE in females but not in males. (h) FH+ females > FH-females in pairwise DMN network TE in transitions to VIS+/- brain states. DAT: (d) The interaction effect was driven by FH+ males < FH-males. (g) Family history density had a significant negative correlation with DAT network TE in males but not females. (i) FH+ males < FH-males in pairwise DAT network TE across almost all transitions. VAT: (e) The interaction effect was driven by FH+ males < FH-males. (h) Family history density had a significant negative correlation with VAT network TE in males but not females. (j) FH+ males < FH-males in pairwise VAT network TE to DMN+/- brain states. *significant before and **significant after correction.

We next investigated which pairwise state transitions drive the observed effects of family history of SUD and sex on mean network TE. Within-sex unpaired *t* tests revealed FH+ females exhibit increased pairwise TE of the DMN across all transitions, particularly transitions to the VIS meta-state, whereas males exhibit no significant differences (Figure 4h). FH+ males have decreased pairwise TE of the DAT for almost all transitions compared to FH-males (Figure 4i) and decreased pairwise TE of the VAT in transitions to the DMN meta-state (Figure 4j), whereas females exhibit no significant differences.

### 2.6 Regional Transition Energy

We next sought to identify regions contributing to observed group-by-sex differences in global and network TE. We ran ANCOVAs on mean regional TE (dependent variable) using the same terms included in the models above. ANCOVA results for all regions can be found in the Supplementary Information. Prior to FDR correction (over the 86 regions), family history of SUD had a significant effect on the regional TE of the bilateral banks of the superior temporal sulcus (banks of STS), bilateral paracentral lobules, bilateral superior temporal gyri, and right amygdala (Figure 5a). The interaction between family history of SUD and sex was significant in the left isthmus cingulate, bilateral pars orbitalis, right supramarginal gyrus, bilateral superior parietal lobules, left pericalcarine, and right cerebellum, prior to FDR correction (Figure 5d).

**Figure 5.**
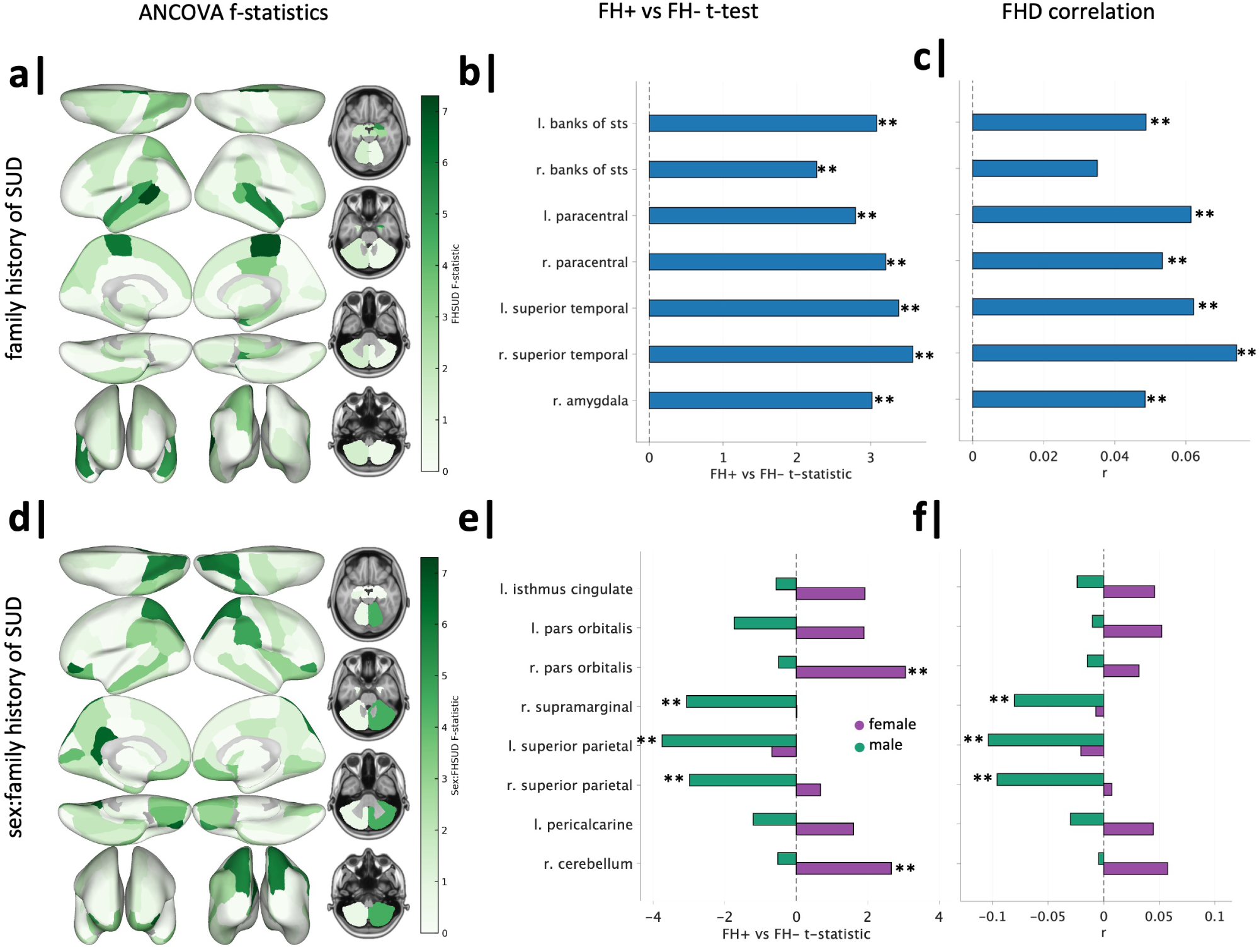
Regional transition energy differences in family history of SUD and family history of SUD by sex. (a) ANCOVA of mean regional TE revealed significant effects of family history of SUD on seven regions, though they did not survive correction. (b) FH+ vs FH-t-tests revealed significant (post-correction across *t* tests of seven regions) FH+ > FH-across all seven regions. (c) Family history density had significant mild positive correlations with regional TE of all seven regions, except the right bank of STS. (d) ANCOVA of mean regional TE revealed significant effects of family history of SUD by sex in eight regions, though they did not survive correction. (e) Within-sex t-tests revealed FH+ females > FH-females in the right pars orbitalis and right cerebellum and FH+ males < FH-males in bilateral superior parietal lobules and right supramarginal gyrus (significant post-correction across *t* tests of eight regions). (f) Family history density had significant but weak negative correlations with regional TE of bilateral superior parietal lobules and right supramarginal in males. *significant before and **significant after correction.

Unpaired post hoc *t* tests were performed to compare FH+ and FH-individuals’ TE for regions found to have a significant effect of family history of SUD and within each sex for regions with significant group-by-sex effects. Across all seven regions analyzed, FH+ individuals (regardless of sex) had significantly increased regional TE compared to FH-individuals (Figure 5b). Further, all regions (except the right bank of STS) were found to have a significant, though weak, positive correlation with family history density of SUD (Figure 5c).

Within-sex *t* tests revealed significant increases in regional TE of the right pars orbitalis and right cerebellum were found in FH+ females compared to FH-females. Bilateral superior parietal lobules and the right supramarginal were significantly decreased in FH+ males compared to FH-males (Figure 5e). The direction and effect size of correlations between family history density and regional TE of these eight regions largely recapitulated the results of within-sex t-tests. The right supramarginal (pre-FDR correction) and bilateral superior parietal lobules (post-FDR correlation) demonstrated significant, yet mild, negative correlations with family history density in males (Figure 5f). Group differences in pairwise regional TE of these regions were consistent with mean regional TE results shown here (Figures S3 & S4).

### 2.7 Robustness Analyses

To ensure the robustness of our results, we replicated our main findings in several ways: (i) with *k*=5 clustering, (ii) with individual structural connectomes (SC) in a cortex-only parcellation, and (iii) in a cohort of sex, age, and in-scanner motion-matched subjects from an external dataset (National Consortium on Alcohol and NeuroDevelopment in Adolescence; NCANDA) (60). We also re-ran ANCOVA models by stratifying our cohort in various ways: (i) a single site with the largest number of subjects, (ii) each MRI scanner model, and (iii) each income category.

We found that our main results were consistent when varying the number of clusters (*k*=5) (Figure S8), and when using individual SC’s (Figure S11). Our independent analysis of the NCANDA dataset showed similar trends of FH+ > FH-females and FH+ < FH-males at the global and regional levels, further supporting the generalizability of our findings across different populations (Figure S9). Subjects within the largest site (site 16) displayed significant family history-by-sex interactions on global TE (FH+ > FH-females and FH+ < FH-males) and in DMN TE (FH+ > FH-females), but not in the DAT or VAT (Figure S12. Analyzing data by MRI scanner model, we observed that our main results were largely consistent in subjects scanned with Siemens models, while results from the GE scanner differed, likely due to demographic differences and lower data quality (Figure S13). See Table S2 for subject demographics by MRI model. Previous ABCD analyses found GE scanners have lower reproducibility in diffusion MRI metrics (61), higher distinguishability after site-normalization in rsfMRI data (62), and higher non-compliance to imaging protocols across modalities (63) compared to Siemens. Further, Siemens scanners implement real-time motion monitoring while GE scanners do not (64); motion correction is critical in this dataset of young adolescents. When stratifying the cohort by income level, our primary findings were consistent mainly in the highest income group (i.e., largest subgroup) (Figure S14). In the ABCD cohort, previous work has found higher income is related to greater curiosity and availability of substances, higher rates of early substance use, and greater odds of substance use in this cohort (65; 66; 67). Overall, these analyses confirm the robustness and reliability of our findings across various conditions and datasets, but indicate a possible influence of socioeconomic and demographic factors.

## 3 Discussion

Using a network control theory approach that models the brain as a networked, complex, dynamical system, we investigated differences in the brain activity dynamics of substance-naïve youth with and without a family history of SUD. Our findings indicate that a family history of SUD was associated with increased TE in certain brain regions, and that the effect of family history varies by an individuals’ sex at all levels of investigation, including global, network, and regional. Specifically, we found females with a family history of SUD tended to have increased TE compared to females without a family history, while the opposite was true for males. Interestingly, increases in FH+ females were driven by the default mode network, while decreases in FH+ males were driven by dorsal and ventral attention networks.

### 3.1 Family history of SUD increases regional transition energies in specific regions, independent of sex

Family history of SUD was associated with increased TE in several regions: bilaterally in the banks of the STS, paracentral lobules and superior temporal gyri, and in the right amygdala. Functional and structural abnormalities of these regions have been observed in individuals with SUD or high risk of SUD (17; 58; 68; 69; 70; 71; 72). A meta-analysis by Pando-Naude et al. (17) found individuals with various types of SUDs consistently exhibit reductions in the volume of the superior temporal gyrus, a region shown to have increased engagement during craving in SUD (73; 74; 75). Studies in substance-naïve FH+ youth from the ABCD baseline assessment have reported reduced cortical thickness in left bank of STS and left paracentral lobule, as well as greater left paracentral activation during successful inhibition (58; 76). Further, family history of alcohol use disorder (AUD) is consistently associated with reduced volume of the amygdala (68; 69; 70), a key region in withdrawal and craving symptoms of SUD (77). We found a significant effect of family history in the right amygdala, but not the left, perhaps reflecting functional lateralization of emotional processing in the amygdalae (78). Taken together, our findings of increased TE in these regions may be the result of structural and/or functional alterations and reflect heightened risk of SUD.

### 3.2 Family history of SUD in females increases regional and default mode network transition energies

FH+ females exhibit higher TE in the DMN, a network linked to self-generated thought, monitoring the internal state, and higher-order cognition (79; 80; 81; 82; 83). Atypical DMN activity has been previously observed in individuals with parental history of SUD (51) and with ongoing SUD (84; 85; 86; 87). Recent models suggest the DMN, positioned at the top of the brain’s network hierarchy (88), plays are role in optimizing prediction errors and exerts top-down inhibitory control over attentional and sensory systems (82; 83; 88). Given that increases in DMN TE of FH+ females are driven by transitions to the VIS meta-state, our findings suggest greater difficulty in carrying out “top-down” transitions theoretically important for inhibitory control and supports the idea that SUD results from disrupted top-down communication (89; 90). These findings provide a mechanistic explanation for FH+ females’ heightened propensity for rapid habit formation (32). Additionally, abnormal DMN activity, linked to withdrawal and craving in SUD (84; 91), may cause rumination on negative internal states, aligning with observations of females’ susceptibility to negative reinforcers of substance use (32; 92).

Regionally, FH+ females exhibit increased TE of the right pars orbitalis and right cerebellum. The pars orbitalis, a region of the DMN involved in inhibition of inappropriate motor responses and regulation of craving (93; 94; 95), has been linked to future substance use through lower gray matter volume in the ABCD cohort (96). Reductions in cerebellar gray matter volume are frequently observed in SUD (97; 98; 99; 100). Moreover, FH+ youth show reduced cerebellar activation during risky decision-making (101). While the majority of studies have not considered group-by-sex effects (93; 94; 95; 97; 98; 99), some studies note female-specific structural and functional alterations of these regions in SUD or a family history of SUD (76; 102; 103).

### 3.3 Family history of SUD in males decreases regional and attentional network transition energies

FH+ males exhibit a distinct pattern of decreased TE in attentional networks, which appears to align with findings of heightened risk of SUD in youth with behavioral attention deficits (104), atypical functional activity in the DAT and VAT of individuals with SUD (19; 105; 106; 107), and a sustained attention network that predicts future substance use (29). Regionally, FH+ males exhibit decreased TE bilaterally in the superior parietal lobules and in the right supramarginal gyrus, which are engaged in goal-directed attention (108) and cognition and language (109), respectively. Altered structural and functional neurophenotypes have been observed in youth from the ABCD cohort with a parental history of AUD (58; 76), where family history-by-sex effects were either not found (76) or not considered (58). Further, a study in substance-naïve youth identified decreased cortical thickness of the left supramarginal gyrus as the top predictor and being male as second in predicting future alcohol use (110). Our findings suggest reduced energetic demand in attention networks and regions prior to substance use may lead an individual to more readily attend to the rewarding effects of substances once exposed. This is particularly evident in FH+ males’ decreased pairwise TE of the VAT, a network responsible for stimulus-driven allocation of attention, being driven by transitions to the DMN+/- brain states, representing “bottom-up” state transitions.

### 3.4 Female and male FH+ youth exhibit distinct halves of a common neurodevelopmental model of SUD

How differences in the resting brain’s transition energies relate to SUD predisposition remains unclear. One possibility is that higher or lower energetic demand of a network/region makes it less or more dominant, respectively, in driving brain state transitions during action and processing of the environment. This theory, based on the assumption of energy consumption minimization (52; 111), coupled with our current findings, suggests a reduced ability of FH+ females to reach states that support impulse inhibition by the DMN, and an increased propensity of FH+ males to occupy states driven by reward saliency via the DAT/VAT. Put another way, FH+ females have a harder time “stepping on the brakes”, whereas FH+ males more readily “step on the gas”. This aligns with a common disease model of addictive behaviors, which suggests a strengthened impulsive/reactive system (bottom-up) and a diminished reflective/deliberative system (top-down) (21). While acute and chronic effects of drugs may have both effects, our findings suggest FH+ females and males premorbidly exhibit distinct halves of this model. Additionally, abnormality in a reflective/deliberative system in females and an impulsive/reactive system in males aligns with the sexes’ respective higher probabilities for internalizing and externalizing pathways to SUD (35; 36; 37; 38).

Baseline sex differences in the functional dynamics of these networks may explain sex-modulated manifestation of familial predisposition to SUD (112; 113). Further, activity of the DMN and DAT are anti-correlated, meaning when one is active, the other is suppressed (88; 114; 115), a phenomenon evident from early infancy and increasing with maturity (116; 117). Given the inverse functional relationship of these networks and the opposite direction of effect in FH+ females and males (increased DMN TE and decreased DAT TE, respectively), predisposition to SUD in females and males may be underpinned by distinct mechanisms with similar behavioral phenotypes.

### 3.5 SUD vulnerability manifests in functional dynamics of the cortex prior to substance use

Notably, the majority of regions and networks implicated here are cortical, with the exception of the cerebellum and amygdala. Our findings contrast a body of work implicating the subcortical dopaminergic system in SUD predisposition (6; 118; 119; 120; 121). However, these studies largely utilized older subjects who had likely initiated substance use (6; 118; 121). Differences in subcortical regions may emerge in late adolescence/early adulthood through developmental processes or in response to the onset of substance use (118; 122). Moreover, our findings echo that of Pando-Naude et al. (17) who propose addiction is characterized by both cortical and subcortical pathologies, whereas occasional use (risk factor for future dependence) is characterized by cortical pathology only. Recent findings applying NCT in young adults with current heavy alcohol use identified increased global TE and, importantly, increased TE in transitions from subcortical to higher-order states (18). Taken together, our results posit cortical abnormalities are present in FH+ youth prior to substance initiation, with perhaps further cortical and subcortical pathologies resulting from active and/or chronic substance use. Furthermore, our findings pinpoint functional alterations as more prominently altered in this cohort of individuals. Using individual structural connectomes, we found the majority of our main results are replicated (S11) and global TE metrics were highly correlated as those from a group-average SC. These findings align with recent observations of a lack of differences in white matter integrity between FH+ and FH-drug-naive adolescents, suggesting familial predisposition to SUD is manifested primarily in functional dynamics (26).

### 3.6 Limitations and Future Directions

The age range of cohort our represents a period of major neurodevelopmental changes, which vary by sex (53; 104; 123; 124; 125; 126). Thus, our findings need to be validated over the entire window of development through leveraging the longitudinal nature of the ABCD and NCANDA studies. Notably, we did find our mean global TE results replicated in the NCANDA dataset in individuals aged 12-22 years old. Further, exclusion of subjects with in-scanner motion may have implications on reported findings given that head motion is a stable and heritable phenotype in adults (127; 128) and a phenotypic marker of impulsivity (129). Increased head movement (even within acceptable limits) was shown to be an important variable in a machine learning study predicting future alcohol use (5). Therefore, our cohort may not represent the full range of extremity of SUD predisposition. Additionally, our analyses did not examine differences by gender identity due to the limited gender diversity in the ABCD baseline assessment. As the cohort matures, examining the effect of sex, gender and sexual orientation on SUD predisposition will be critical, particularly due to higher rates of SUD in lesbian, gay, bisexual, transgender, and queer (LGBTQ+) youth (130; 131; 132).

## 4 Conclusion

Our study reveals sex-specific effects of a family history of SUD with distinct network alterations in male and female youths with a family history: FH+ males exhibit decreased transition energy in the attentional networks and FH+ females show heightened transition energy in the DMN. We posit that this may translate to FH+ males more readily “stepping on the gas” and FH+ females having a harder time “stepping on the brakes” in terms of substance use patterns. These findings suggest that mechanisms underlying SUD predisposition are complex and influenced by sex-specific neurodevelopmental trajectories that may ultimately result in a common constellation of behavioral phenotypes. Recognizing these mechanistic differences is crucial for understanding the onset of SUD and developing targeted intervention strategies.

## 5 Methods

### 5.1 Sample Characteristics

The Adolescent Brain and Cognitive Development (ABCD) study is longitudinally tracking the brain development and health of a nationally representative sample of children aged 9 to 11 years old (at the time of enrollment) from 21 centers across the United States (https://abcdstudy.org). Parents’ full written informed consent and all children’s assent were obtained by each center. Research protocols were approved by the institutional review board of the University of California, San Diego (no. 160091), and the institutional review boards of the 21 data collection sites (133). The current study utilized neuroimaging data from the 2.0.1 release and non-imaging instruments from the baseline assessment updated to ABCD Data Release 5.1.

#### 5.1.1 Exclusions

From the original ABCD cohort of N=11,868, we excluded youth who: i) did not survive strict MRI quality control and/or exclusion criteria as established by Chen et al. (134) (N = 9506), ii) were scanned on Philips scanners (N = 2), iii) did not meet criteria for group definitions of FH+ or FH- (see Section 5.1.2; N = 109), iv) had previously used substances (see Section 5.1.3; N = 54), v) were adopted (N = 32), vi) had a mismatch between reported sex and their sex determined by salivary samples (N = 20), vii) had missing household income information (see Section 5.1.4, N = 119), or viii) had missing information on parental mental health issues (5.1.2, N = 79). Further, 53 subjects were excluded (after *k*-means clustering and TE calculations) due to outlier mean global TE values (see Section 5.3.3). See Table S1 for excluded subject demographics.

#### 5.1.2 Family History of Substance Use Disorder and Mental Illness

We utilized the baseline assessment of the Family History Module Screener (FHAM-S) (135; 136) in which parents reported psychopathology and substance use problems within the first and second degree biological family members of each child. Although family history of alcohol and drug problems were obtained separately, in analyses we considered cross-substance family history of substance use (FH+) as having family members with a history of either alcohol or drug use problems. This choice is supported by recent evidence for the heritability of a general vulnerability factor for cross-substance addiction (3) and neuroimaging findings of abnormalities across SUDs to a common brain network that is similar across imaging modalities and substances (137). In the FHAM-S, problems with drugs or alcohol may include marital, work, school, arrests or driving under influence, health, rehabilitation programs, heavy use, or social issues.

Following previous work (56; 57; 58), we categorize individuals as FH+ if they have at least one parent or a minimum of two grandparents with a history of SUD. Individuals with no parents or grandparents with a history of SUD are classified as family history negative (FH-). Individuals with just one grandparent with a history of SUD are classified as FH+/- and are only included in analyses that rely on continuous associations (i.e., family history density) as these individuals likely have minimal genetic load of previous generations (56), yet could not be classified as FH-.

In addition to FH+ or FH-group assignments, a more granular measure of family history density (FHD) was calculated based on the sum of positive reports of problems from biological parents (+1) and biological grandparents (+0.5). FHD scores could thus range from 0 to 4, with a score of 0 indicating no family history of SUD and a score or 4 indicating SUD in both parents and all four grandparents (58). Participants with only one grandparent with a history of SUD, who were excluded from categorical analyses (FH+ vs FH-), were included in FHD analyses. As in previous work (57; 58), other second-degree relatives (i.e., aunts and uncles) were not considered due to missingness and less accurate account of distal relatives.

Given the high comorbidity of substance use problems and mental illness, we also included a binary variable for parental history of mental illness, other than SUD, as a variable in our analyses. The FHAM-S evaluates parental history of mental health issues including: suicide, depression, mania, anti-social personality, schizophrenia, and other emotional/mental heath issues. If one or more parents of a subject were reported as experiencing one or more of these mental health issues, a subject met criteria for having a parental history of mental health issues.

#### 5.1.3 Exposure to Substances

##### Childhood substance use

To isolate the effects of family history of SUD from substance use itself, we excluded youth who had initiated substance use according to parents or children themselves (N = 54). Subjects were excluded if they self-reported lifetime use of more than a sip of alcohol, more than a puff of a cigarette/e-cigarettes, or any use of nicotine products, cannabis products, synthetic cannabinoids, cocaine, cathinones, methamphetamine, ecstasy/MDMA, ketamine, gamma-hydroxybutyrate, heroin, psilocybin, salvia, other hallucinogens, anabolic steroids, inhalants, or prescription misuse of stimulants, sedatives, opioid pain relievers or over-the-counter cough/cold medicine (138). Parents were also asked if their child had exhibited drinking problems (consumed three or more drinks a day, consumed two drinks in the last 12 months) or used drugs (cocaine, marijuana, solvents, stimulants, tobacco, opioids, hallucinogens, sedatives, or other). If parents endorsed any of these, children were excluded from analyses.

##### In utero substance exposure

Given the known impact of maternal substance use during pregnancy on brain development (139; 140), we included in utero substance exposure as a binary variable in our ANCOVA models. Prenatal substance use exposure was reported based on caregiver recall in the ABCD Developmental History Questionnaire (136). Consistent with previous work using dichotomous analyses (141), we considered exposure as either present or absent based on if mothers reported maternal use of over 7 drinks of alcohol per week (142), any tobacco, marijuana, cocaine, opiates, or other drugs after pregnancy was recognized.

#### 5.1.4 Household Income and Parental Education

We included two measures typically associated with socio-economic status (SES): household income (HI) and parental education (PE). Both of these variables are considered essential for mental health research by the National Institute of Mental Health (143) and have a known relation to substance use disorders and their consequences (144; 145). In the ABCD study, both HI and PE were obtained in parent-reported demographic questionnaires (136). In order to simplify our statistical models without losing nuance, we followed previous work and re-coded the PE and HI variables into fewer groups (146; 147). The PE survey responses, for which there were originally 21 choices, was re-coded into 5 levels: less than high school, high school graduate/GED, some college, associate’s/bachelor’s degree, and postgraduate degree. We used the highest of the two parents PE. For HI, the original nine response options were re-coded into three levels: (1) less than $50,000, (2) more than $50,000 and less than $100,000, and (3) over $100,000. Individuals missing PE or HI data were excluded from analyses.

#### 5.1.5 Sex Assigned at Birth

Our analysis of sex differences focuses on sex assigned at birth, which we refer to as “sex”. We did not control for or assess gender-based differences in brain dynamics in the present study. We excluded individuals whose sex assigned at birth did not match their sex as determined in a salivary sample as this could imply clerical errors in reporting sex assigned at birth. However, we did not control for or assess gender-based differences in the present study. While “male” and “female” brain features exist in both sexes, some features may be more prevalent in one sex than the other (123; 148; 149) and, further, recent work has suggested sex and gender may be mapped on to different brain networks (150). In our current study we have found sex to be a useful, easily obtained, and important biomarker as we have identified meaningful variance within and between each sex.

### 5.2 Neuroimaging Data

#### 5.2.1 Parcellation

In our main results, we present analyses in which the rsfMRI timeseries and group-average structrual connectome (SC) were parcellated using an 86-region atlas derived from FreeSurfer (FS86) combining the 68 region Desikan-Killiany (DK68) gyral atlas (34 cortical regions per hemisphere) with 16 subcortical structures (8 per hemisphere, excluding brainstem) and 2 cerebellar structures (1 per hemisphere) to render a whole brain anatomically-defined parcellation for each subject (151; 152). Each cortical region was assigned to one of seven networks from the functionally-defined Yeo seven-network parcellation (59). Subcortical regions were assigned to a subcortical network and cerebellar regions to a cerebellar network. The rsfMRI time-series and individual structural connectome (SC) were also extracted in the DK68 cortical atlas. See Section 5.2.3 for more information on atlas choice.

#### 5.2.2 Resting-state Functional Magnetic Resonance Imaging (rs-fMRI)

We analyzed rsfMRI collected during the baseline assessment of the multi-site ABCD study. Minimally processed rsfMRI data underwent pre-processing and quality control as described in (134; 153) and summarized here. Subjects whose scans were obtained on Philips scanners were excluded due to incorrect post-processing per recommendations of ABCD Consortium. Scans underwent removal of initial frames and alignment with the T1 images using boundary-based registration (BBR). Volumes (as well as one volume before and two volumes after) were marked as outliers if they had framewise displacement (FD) > 0.3 mm or DVARS > 50. Uncensored segments of data containing fewer than five contiguous volumes were also censored. Functional runs were excluded if over half of the volumes were censored or BBR costs were >0.6. Nuisance covariates (global signal, motion correction parameters, average ventricular signal, average white matter signal, and their temporal derivatives) were regressed out of the fMRI time-series using non-censored volumes to estimate regression coefficients. Brain scans were band-pass filtered (0.009 *≤ f ≤* 0.08 Hz), projected onto FreeSurfer fsaverage6 surface space, and smoothed using a 6-mm full-width half-maximum kernel. Upon receiving this pre-processed data, we subsequently removed censored volumes from the time-series and normalized the BOLD time series by the mean gray matter BOLD signal (pre-spatial and temporal filtering). After censoring outlier frames, the average number of remaining rsfMRI frames (across all scans) was 1155.88±291.12 (mean ± std).

Given the known scanner effects in the rs-fMRI data of the ABCD dataset (62), we included the scanner model in all ANCOVA models as a covariate of no interest. As Philips scanners were excluded, our dataset consisted of three scanner model types: Prisma (Siemens), Prisma Fit (Siemens), and Discovery MR750 (GE).

#### 5.2.3 Structural Connectomes

In the ABCD study, diffusion MRI was collected at the baseline assessment (154). Pre-processed dMRI was further processed by deterministic tractography with SIFT2 global streamline weighting and regional volume normalization. Structural connectomes (SC) were extracted in the FS86 (cortical and subcortical) for 149 subjects and DK68 (cortical only) for 2080 subjects. The SC matrices are symmetric with the diagonal (self-connections) set equal to zero. Global TE values using a group-average SC and individual SC for N subjects were found to be highly correlated (rho = 0.99, *p* < 0.0001). Given this high correlation and the known relevance of subcortical regions in SUD literature (6; 118; 119; 120; 121), we chose to utilize the group-average SC for the FS86 atlas. This choice is further supported by previous NCT work (50) and recent observations of a lack of differences in white matter integrity between FH+ and FH-drug-naive adolescents, suggesting familial predisposition is manifested primarily in functional dynamics (26). Main results were replicated in the DK68 parcellation using individual, cortex-only SCs (**Figure S11**).

#### 5.2.4 Framewise Displacement

Given the concern for head motion during neuroimaging of a pediatric cohort, we controlled for subjects’ tendency to move their head during rest in the scanner by calculating the average framewise displacement (FD) across all timepoints of all scans per subject (censored and uncensored). We included mean FD as a covariate of no interest in all ANCOVA models.

### 5.3 Network Control Theory Analyses

#### 5.3.1 Extraction of brain states

Following Cornblath et al. (47), all subjects’ fMRI time series were concatenated in time and *k*-means clustering was applied to identify clusters of brain activation patterns, or brain states. Pearson correlation was used as the distance metric and clustering was repeated 10 times with random initializations before choosing the solution with the best separation of the data. To further assess the stability of clustering and ensure our partitions were reliable, we independently repeated this process 10 times and compared the adjusted mutual information (AMI) between each of the 10 resulting partitions. The partition which shared the greatest total AMI with all other partitions was selected for further analysis. In general, we found that the mutual information shared between partitions was very high (>0.99), suggesting consistent clustering across independent runs. We chose the number of clusters textk via the elbow criterion, i.e. by plotting the gain in explained explained across clusterings for *k*=2 through *k*=14 and identifying the “elbow” of the plot, which was at *k*=4 (Figure S1). In addition, *k*=4 fell below 1% of variance explained by clustering, a threshold used previously for determining *k* (47; 55). Thus, we chose *k*=4 for its straightforward and symmetric interpretation and replicated main results for *k*=5 in Supplementary Information. For interpretability, each cluster centroid was named via one of nine a priori defined canonical resting-state networks (RSN) (59) plus subcortical and cerebellar networks, by the cosine similarity between the centroid and binary representations of each RSN. Because the mean signal from each scan’s BOLD time series was removed during bandpass filtering, positive values in the centroid reflect activation above the mean (high-amplitude) and negative values reflect activation below the mean (low-amplitude). Subject-specific brain state centroids were calculated for each individual across all included timepoints of their rsfMRI scans.

#### 5.3.2 Transition Energy Calculations

Following similar procedures as described elsewhere (47; 55), we summarize briefly here. We employ a linear time-invariant model:

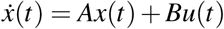

where *A* is a representative (group average) NxN structural connectivity matrix obtained as described above using deterministic tractography from a subset of ABCD subjects (see Section 3.2.3). *A* is normalized by its maximum eigenvalue plus 1 and subtracted by the identity matrix to create a continuous system. *x*(*t*) is a vector of length N containing the regional activation at time t. *B* is an NxN matrix of control points, in this case,the identity matrix is used for uniform control. *u*(*t*) is the external input into the system. N is the number of regions in our parcellation, where N = 86 for our main results. We selected a time-horizon of T = 1.501 as it had the strongest negative correlation with state-pair transition probability measures (across range T = 0.001 to 10), or the likelihood of a transition to occur, as done in previous studies (47; 55). To compute the minimum control energy required to drive the system from an initial brain state to a target state over T, we compute an invertible controllability Gramian for controlling the network A from N nodes.

Using the above methodology to define brain states (see Section 3.3.1), we calculate the regional transition energy for a given transition between each pair of brain states and persistence within each state (pairwise regional TE). To calculate pairwise regional TE, we integrate u(t) over the time-horizon to yield the total amount of input signals into each region necessary to complete each transition, resulting in a *k*x*k* matrix for each region (where *k*= 4 in our main results). We calculate the global transition energy (pairwise global TE), a *k*x*k* matrix, by summing all pairwise regional TE for the transition. We calculate the transition energy required of a network (pairwise network TE) by summing all pairwise regional TE for those regions assigned to a network, resulting in a *k*x*k* matrix for each network. To calculate mean global TE (1 constant), mean network TE (vector of length equal to the number of networks) and mean regional TE (vector of length equal to number of regions), we average across all pairwise TEs at each respective level of analysis.

#### 5.3.3 Defining Outlier Subjects

After calculating mean global TE values, we excluded N = 53 individuals from further analyses due to a mean global TE more than 3 scaled median absolute deviations (MAD) from the sample median (155). While within acceptable limits, these subjects were found to have significantly higher mean framewise displacement compared to non-outlier subjects (*t* = 3.58, *p* = 3.5e-04). Sample characteristics of all excluded subjects can be found in Supplementary Table S1. Importantly, the reported results are largely similar when including outlier subjects.

### 5.4 Statistics

Between-group comparisons were made using ANCOVA models including age, sex assigned at birth (“sex”), family history of SUD, race/ethnicity, parental education, household income, in utero exposure to substances, parental mental health issues, MRI scanner model, in-scanner motion (i.e., mean framewise displacement), and two interaction terms: family history of SUD:sex, and family history of SUD:household income. Full ANCOVA results can be found in the Supplementary Information. For metrics (global, network and regional TEs) found to be significant in ANCOVA models, unpaired *t* tests were performed post hoc between FH+ and FH-subjects across all subjects (significant for family history of SUD) and within each sex separately (significant for interaction between family history of SUD and sex) to determine the direction of effect. Correlations between average TE and family history density were calculated using Spearman’s rank correlations. *P*-values were corrected for multiple comparisons using the Benjamini-Hochberg False Discovery Rate (*q*=0.05) procedure where indicated (pFDR) (156).

## 6 Acknowledgements

We would like to thank Thomas Yeo’s lab, specifically Leon Qi Rong Ooi, for providing the processed MRI data. LS was supported by the National Institute on Drug Abuse of the National Institutes of Health under Award Number T32 285 DA03980. KMP was supported by R01 DA057567 and U24 AA021697. CT was supported with a Career Transition Award (TA-2204-39428) and a postdoctoral fellowship (FG-2008-36976) from the National Multiple Sclerosis Society. AK was supported by the National Institute of Mental Health of the National Institutes of Health under Award Number RF1 MH123232.

## 7 Data Availability

Data used in the preparation of this article were obtained from the Adolescent Brain Cognitive DevelopmentSM (ABCD) Study (https://abcdstudy.org), held in the NIMH Data Archive (NDA). This is a multisite, longitudinal study designed to recruit more than 10,000 children age 9-10 and follow them over 10 years into early adulthood. The ABCD Study® is supported by the National Institutes of Health and additional federal partners under award numbers U01DA041048, U01DA050989, U01DA051016, U01DA041022, U01DA051018, U01DA051037, U01DA050987, U01DA041174, U01DA041106, U01DA041117, U01DA041028, U01DA041134, U01DA050988, U01DA051039, U01DA041156, U01DA041025, U01DA041120, U01DA051038, U01DA041148, U01DA041093, U01DA041089, U24DA041123, U24DA041147. A full list of supporters is available at https://abcdstudy.org/federal-partners.html. A listing of participating sites and a complete listing of the study investigators can be found at https://abcdstudy.org/consortium_members/. ABCD consortium investigators designed and implemented the study and/or provided data but did not necessarily participate in the analysis or writing of this report. This manuscript reflects the views of the authors and may not reflect the opinions or views of the NIH or ABCD consortium investigators. The ABCD data repository grows and changes over time. Processed neuroimaging data by (153) used in this study were uploaded by the original authors to the NDA. Researchers with access to the ABCD data will be able to download the data: https://nda.nih.gov/study.html?id=1368. The ABCD imaging data used in this report came from https://doi.org/10.15154/1504041 and non-imaging data was from the 5.1 release (http://dx.doi.org/10.15154/z563-zd24). These data were used in the analyses described in https://doi.org/10.15154/c9z7-ng36. The ABCD data are publicly available via the NIMH Data Archive (NDA).

Collection and distribution of the NCANDA data were supported by NIH funding AA021681, AA021690, AA021691, AA021692, AA021695, AA021696, AA021697. Researchers with access to the NCANDA data will be able to download the data via https://nda.nih.gov/study.html?id=4513.

## 8 Supplementary Information

### S0.1 Subject exclusions

**Table S1.** Demographic and characteristic data for the full cohort of subjects for whom we received rsfMRI data (134; 153), excluded subjects (via exclusion criteria and outlier global TE values) and all included subjects in our analyses.

|  | Chen et al., 2022 cohort (N = 2362) | Excluded Subjects (N = 468) | Included Subjects (N = 1894) |
| --- | --- | --- | --- |
| **Biological Sex, N(%)** |  |  |  |
| Female | 1265 (54) | 247 (53) | 876 (46) |
| Male | 1096 (46) | 220 (47) | 1018 (54) |
| **Age in months, Mean (SD)** |  |  |  |
| Female | 119.9 (7.50) | 120.08 (7.74) | 120.50 (7.51) |
| Male | 120.42 (7.50) | 120.11 (7.44) | 120.01 (7.44) |
| **Family History of SUD N(%)** |  |  |  |
| FH+ | 534 (23) | 83 (18) | 451 (24) |
| FH- | 1400 (59) | 231 (49) | 1169 (62) |
| FH+/- | 319 (14) | 45 (9) | 274 (14) |
| Missing | 109 (5) | 109 (23) | 0 |
| **Frame-wise Displacement, Mean (SD)** |  |  |  |
| Female | 0.11 (0.07) | 0.12 (0.07) | 0.12 (0.08) |
| Male | 0.12 (0.08) | 0.13 (0.81) | 0.11 (0.07) |
| **MRI Scanner Model N(%)** |  |  |  |
| GE Discovery MR750 | 670 (28) | 142 (30) | 528 (28) |
| Siemens Prisma | 823 (35) | 132 (28) | 691 (36) |
| Siemens Prisma Fit | 867 (37) | 192 (41) | 675 (36) |
| Undefined | 2 (0.08) | 2 (0.4) | 0 |
| **Household Income N(%)** |  |  |  |
| < \$50,000 | 500 (21) | 79 (17) | 421 (22) |
| \$50,000 - \$100,000 | 666 (28) | 97 (21) | 569 (30) |
| \$100,000+ | 1196 (51) | 292 (62) | 904 (48) |
| **Parent Education N(%)** |  |  |  |
| > High School | 68 (3) | 19 (4) | 49 (3) |
| High School/GED | 160 (7) | 33 (7) | 127 (7) |
| Some College | 251 (11) | 67 (14) | 284 (10) |
| Associates/Bachelor | 1207 (51) | 226 (48) | 981 (52) |
| Postgraduate | 673 (28) | 120 (26) | 553 (29) |
| **Race/Ethnicity N(%)** |  |  |  |
| White | 1412 (60) | 218 (47) | 1104 (63) |
| Black | 203 (9) | 62 (13) | 141 (7) |
| Hispanic/Latinx | 447 (18) | 106 (23) | 341 (18) |
| Asian | 58 (3) | 20 (4) | 38 (0.2) |
| Other | 241 (10) | 61 (13) | 180 (10) |
| **In Utero Substance N(%)** |  |  |  |
| Yes | 102 (4) | 29 (6) | 73(4) |
| No | 2260 (96) | 439 (94) | 1821 (96) |
| **Parent Mental Health N(%)** |  |  |  |
| Yes | 1208 (51) | 213 (45) | 995 (53) |
| No | 1034 (44) | 135 (29) | 899 (48) |

### S0.2 Choices in *k* -means clustering

#### S0.2.1 Choosing k

We performed 10 repetitions of k-means clustering for *k*=2 to *k*=14. We quantified the variance explained by clustering as the ratio of between-cluster variance to total variance in the data (47; 55; 157). We chose *k* by plotting the variance gained by increasing *k*, to observe where increasing *k* begins to provide diminishing returns in terms of variance explained. The elbow of the variance explained curve is *k*=4 to *k*=5. We also see that the gain in variance explained by increasing *k* to 4, the variance gained by increasing *k* falls below 1%, the threshold used previously for choosing k (Figure 1b) (47; 55). We therefore analyzed the results with k=4 and 5 and chose to include *k*=4 in the main analysis for ease of interpretation.

**Figure S1.**
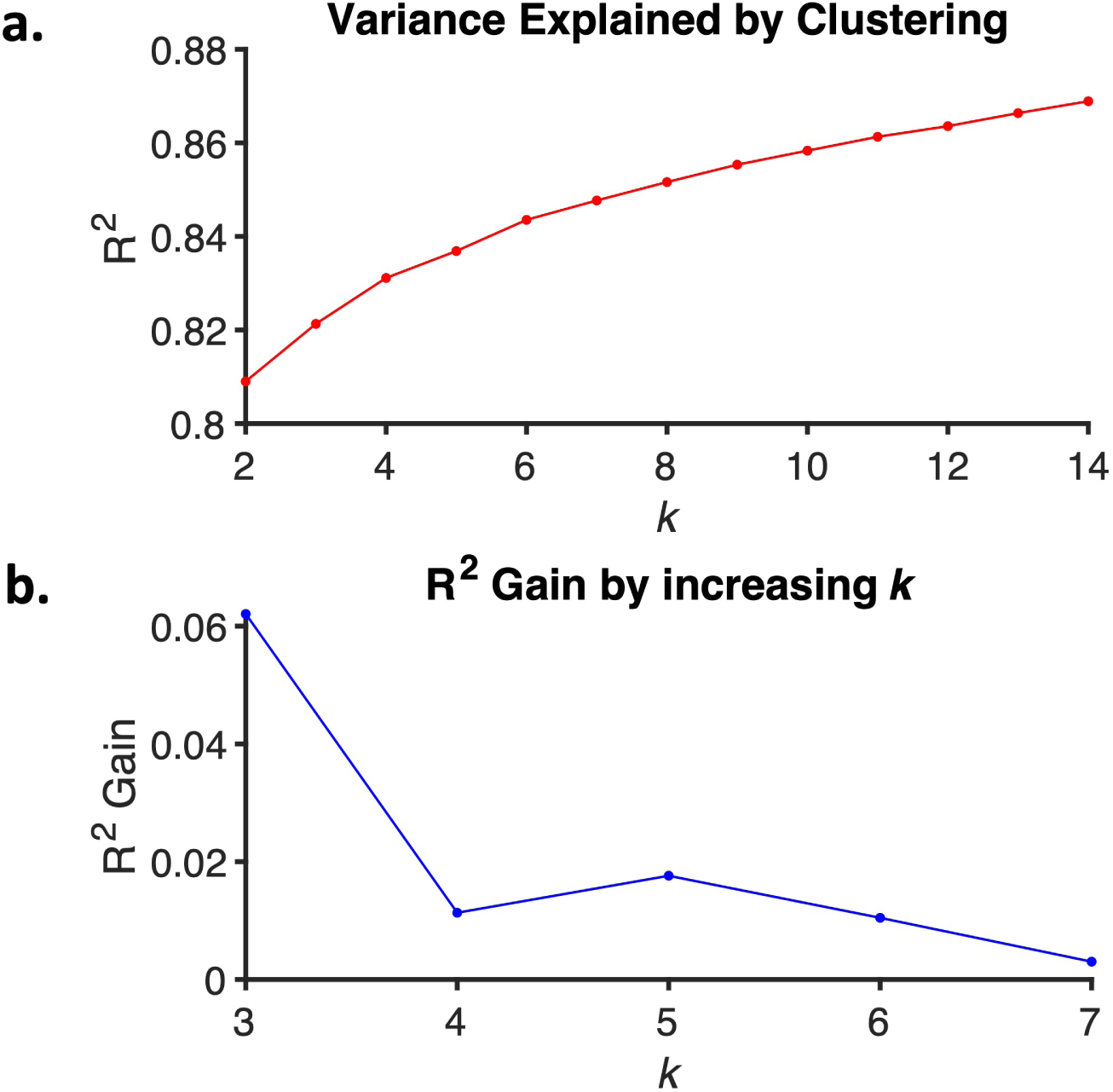
(a) Plot of the variance explained by clustering for each choice of *k*, showing an ‘elbow’ around 4-5. (b) Plot of the gained variance explained by increasing *k*.

#### S0.2.2 Assessing the stability of clustering

As mentioned in the main text, we performed 10 repetitions of k-means clustering, choosing the lowest error solution. To ensure this solution was a consistent global minimum, we repeated this entire process 10 times and compared the adjusted mutual information (AMI) shared between the 10 partitions. The partition that had the maximum sum of AMI scores with all other partitions was selected for analysis. More importantly, this process confirmed that k-means clustering was highly consistent and stable (AMI > 0.99).

**Figure S2.**
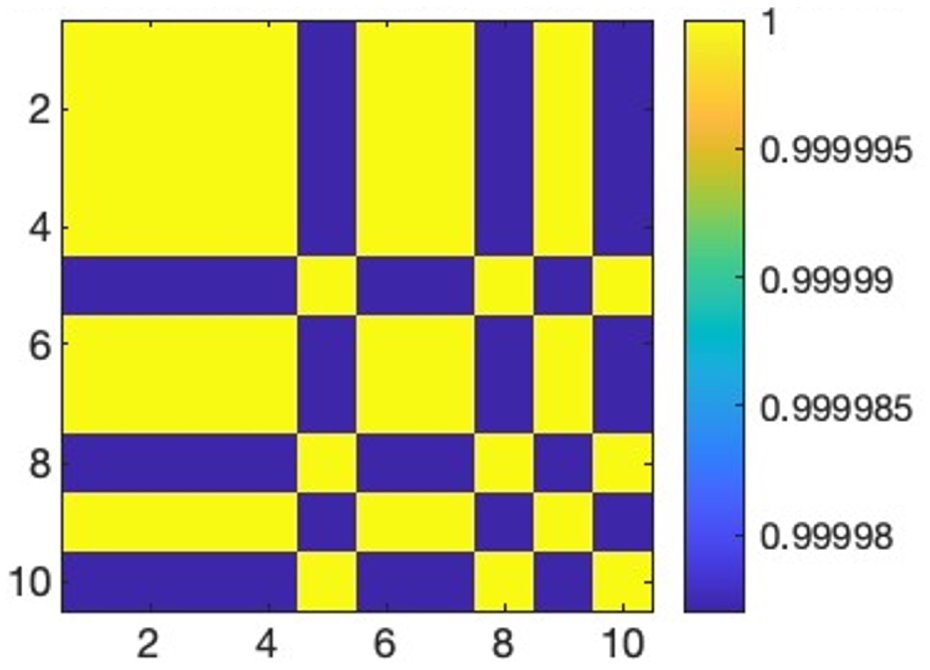
The adjusted mutual information (AMI) shared between 10 independently generated partitions of our data at *k*=4. Values can range from 0 to 1, with 1 indicating identical partitions. The AMI between all partitions was >0.99

### S0.3 Pairwise regional transition energy: post hoc *t* tests

#### S0.3.1 Post-hoc t-tests of pairwise regional transition energy in regions found to have significant effect of family history of SUD (across both sexes)

For each pair of brain states (*k*x*k*), we performed post hoc *t* tests for the pairwise regional TE in each of the seven regions found to be significant for family history of SUD (FH+ vs FH-).

**Figure S3.**
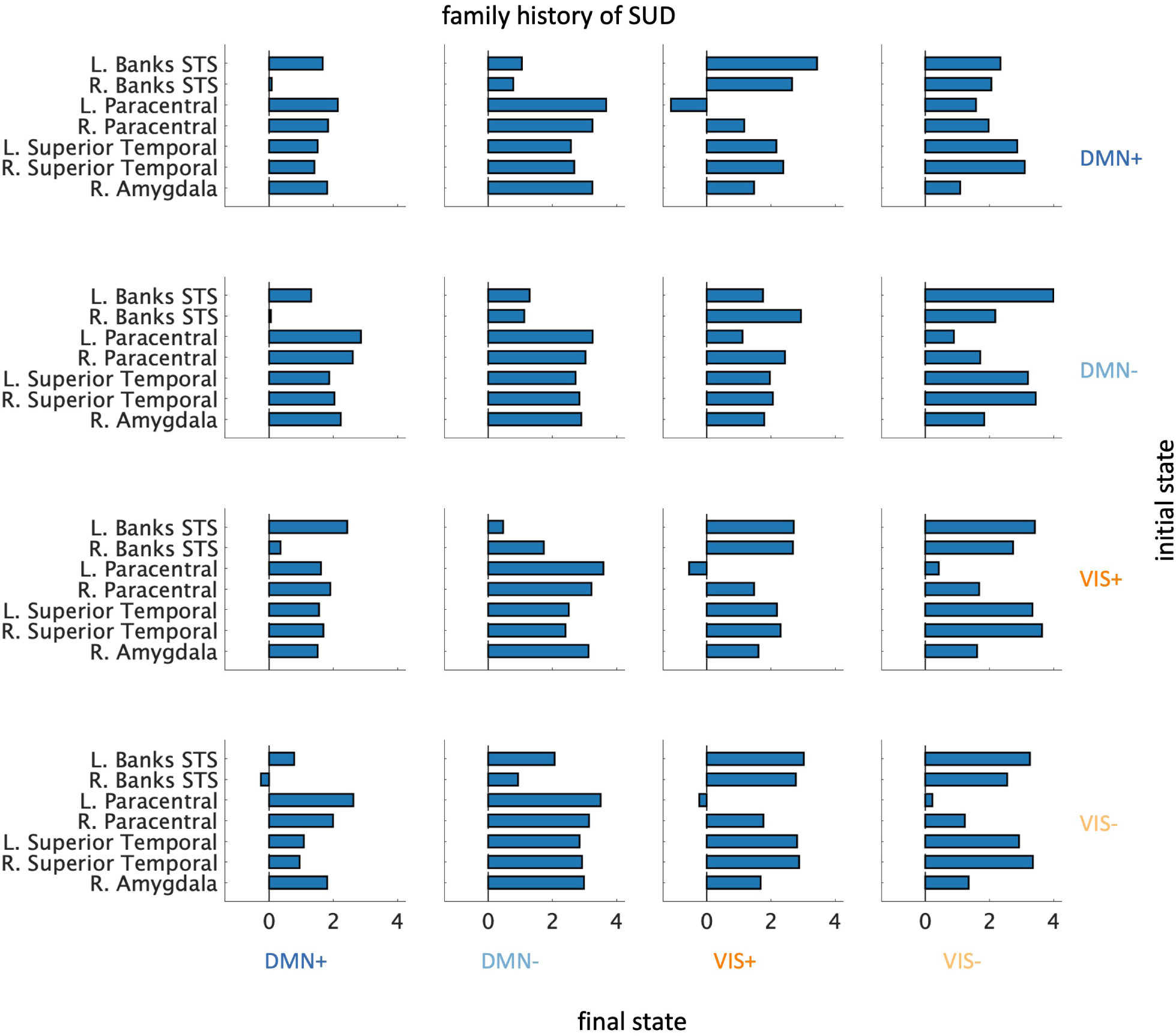
*T* test results for pairwise regional TE values for regions found to have a significant effect of family history of SUD.

#### S0.3.2 Post-hoc t-tests of pairwise regional transition energy in regions found to have significant effect of the interaction between sex and family history of SUD

For each pair of brain states, we performed post hoc *t* tests for the pairwise regional TE in each of the eight regions found to be significant for the interaction of sex and family history of SUD (within-sex FH+ vs FH-).

**Figure S4.**
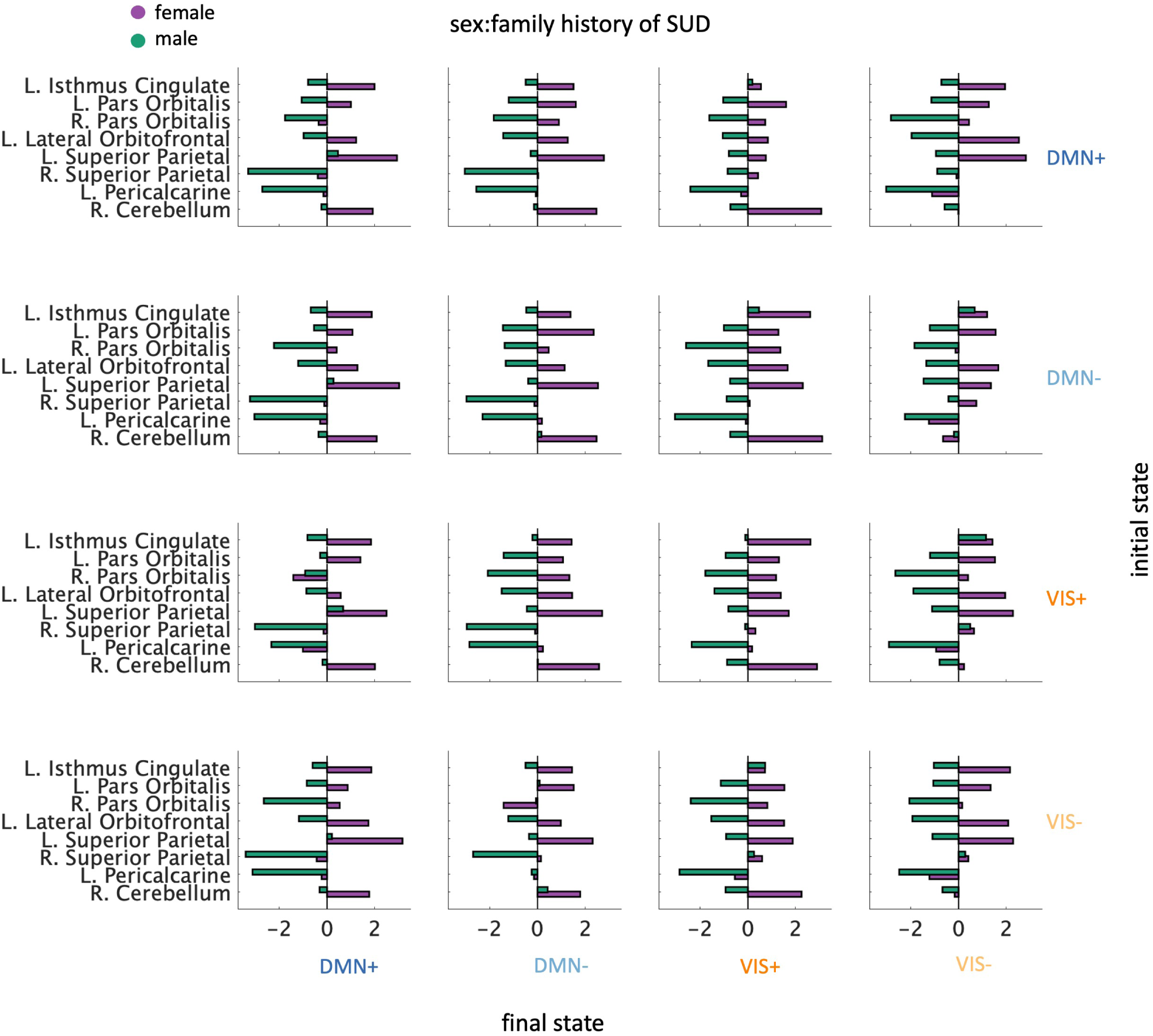
Within sex *t* test results for pairwise regional TE values for regions found to have a significant family history-by-sex effects.

### S0.4 ANCOVA F Statistics for all included covariates

#### S0.4.1 Network TE: ANCOVA F Statistics for all included covariates

**Figure S5.**
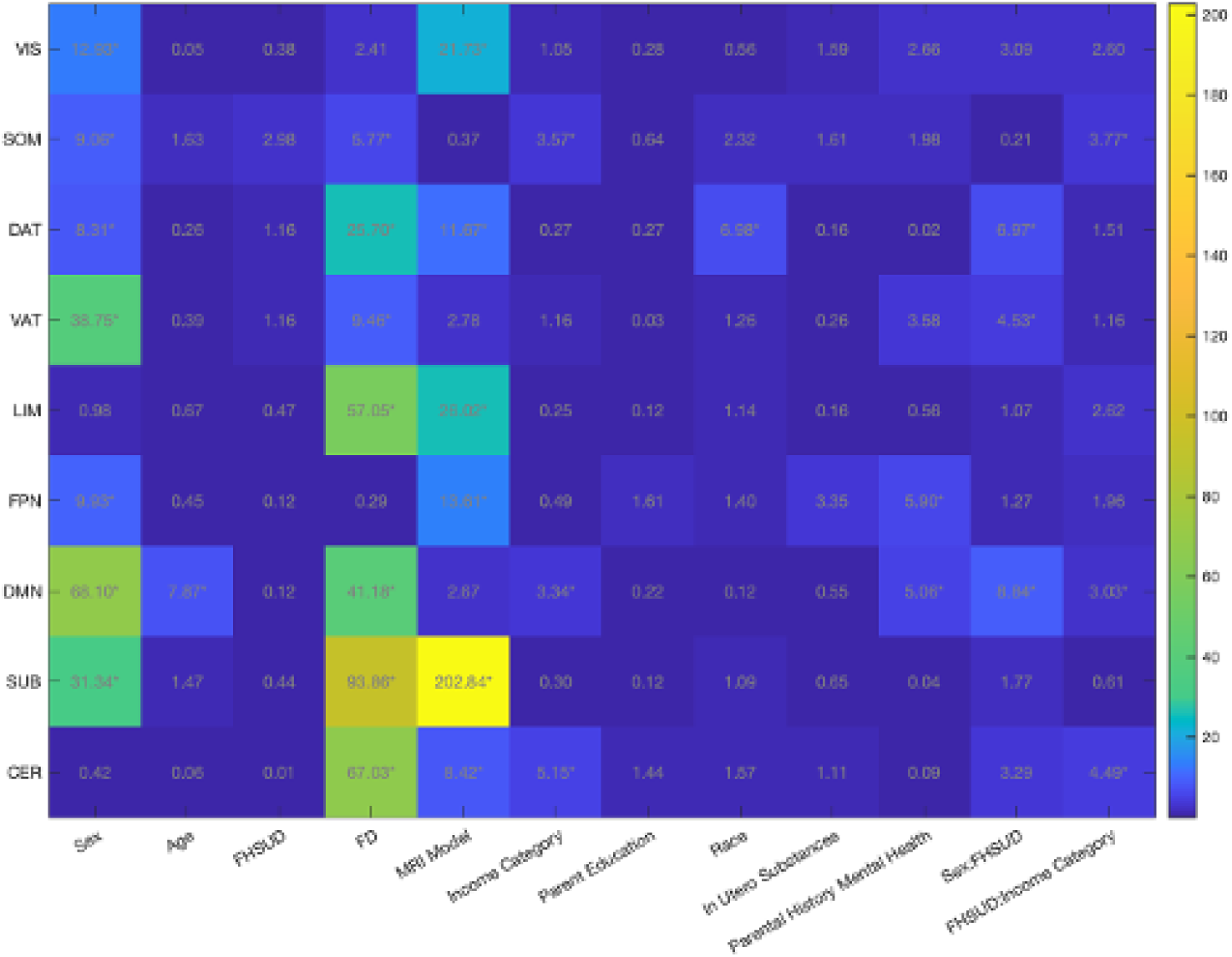
ANCOVA F-statistics for mean network TE values for all variables included in model.

#### S0.4.2 Regional TE: ANCOVA F Statistics for all included covariates

**Figure S6.**
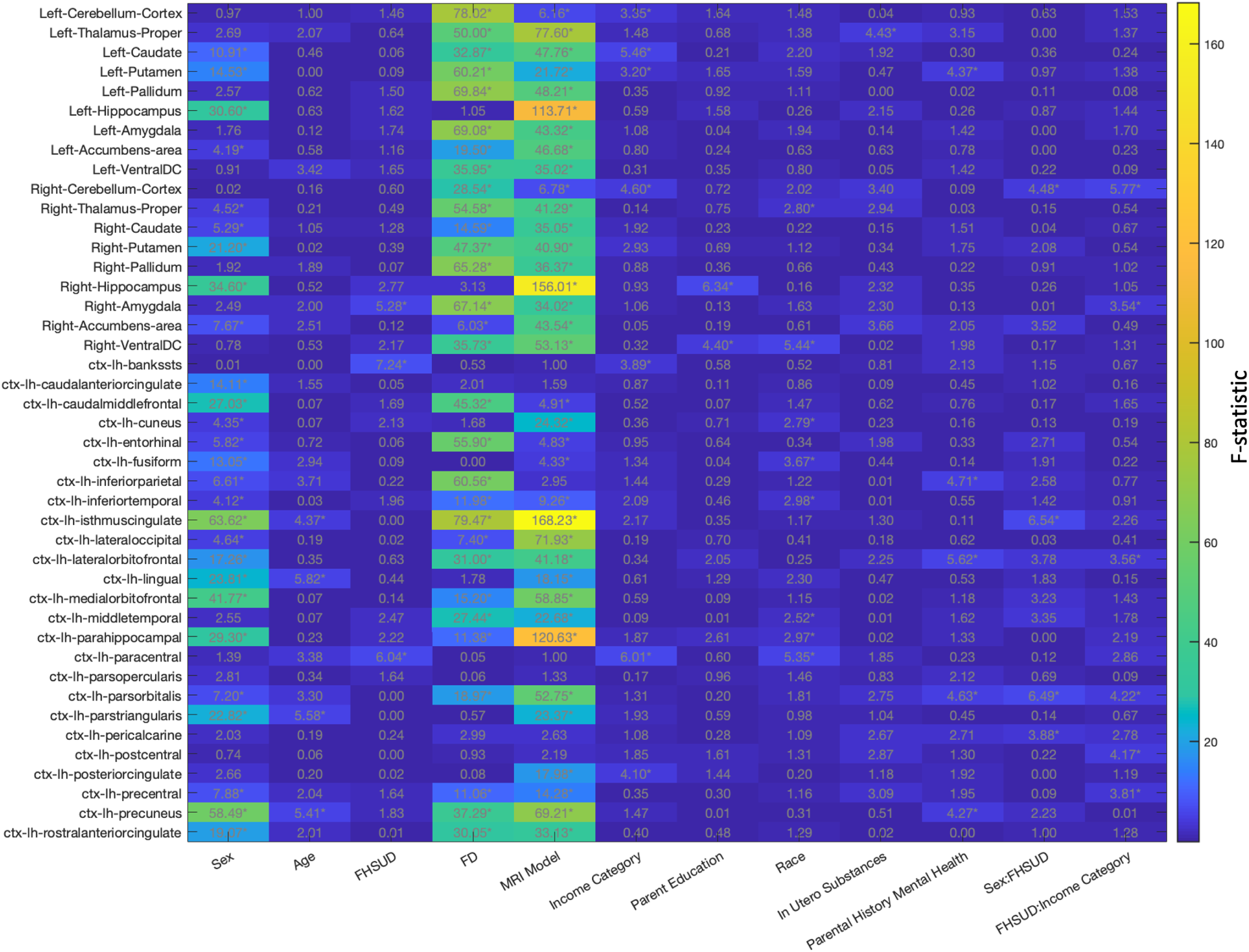
ANCOVA F-statistics for mean regional TE values for all variables included in model. Regions 1 to 43.

**Figure S7.**
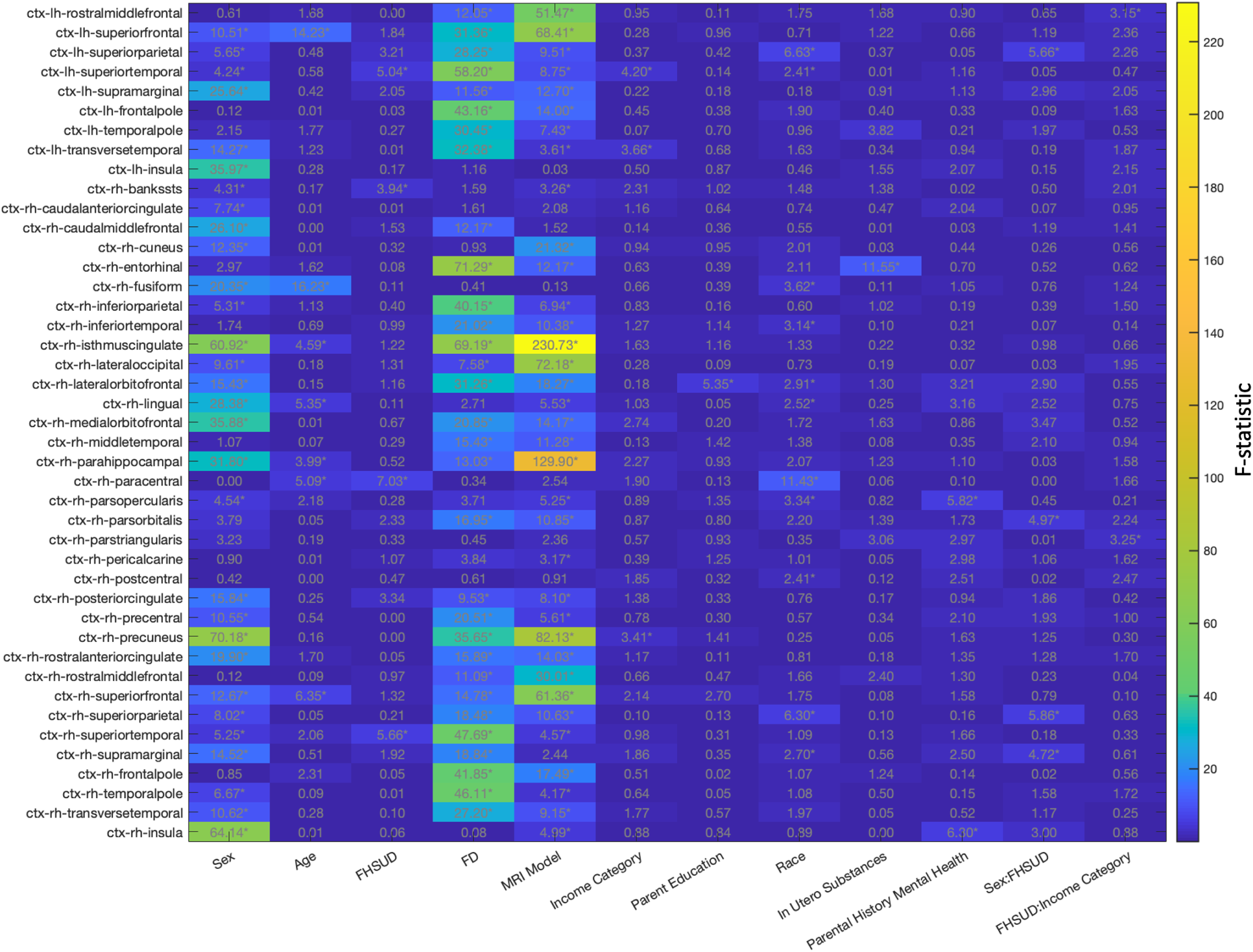
ANCOVA F-statistics for mean regional TE values for all variables included in model. Regions 44 to 86.

### S0.5 Robustness analyses: replications

#### S0.5.1 Replication of main results with k = 5

We replicate our main analysis with *k*=5 to demonstrate the robustness of our results. At *k*=5, the same four brain states as found in *k*=4 plus a state defined by positive amplitude activity in the limbic network (LIM+) were identified as brain states. All major results and trends hold.

**Figure S8.**
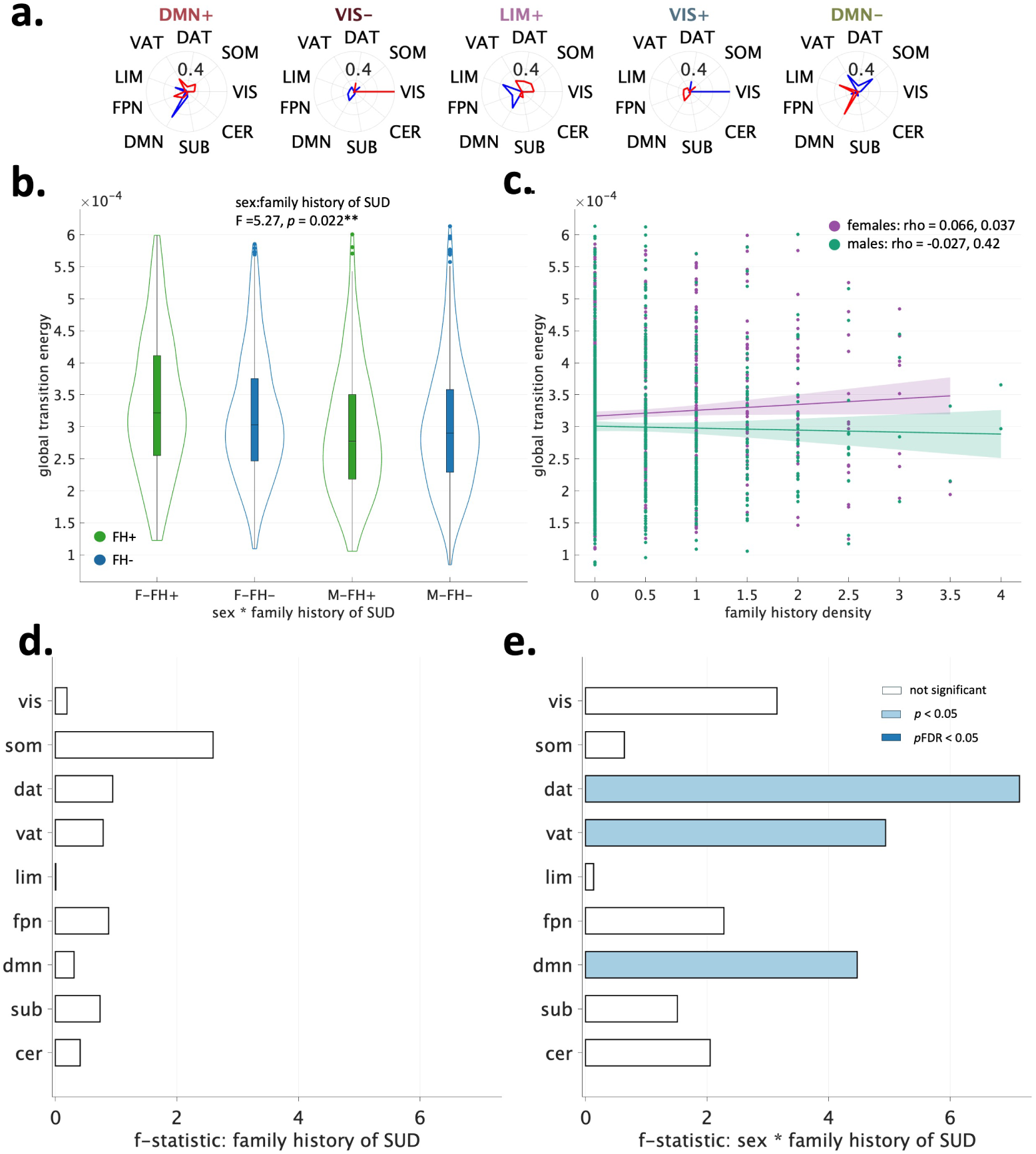
Replication of main analysis with k=5. For b-c, * = *i*<0.05, ** = *p*FDR < 0.5.

#### S0.5.2 Replication of global TE results in adolescents and young adults from the NCANDA dataset

We replicated our results in a cohort of sex, age and in-scanner motion matched individuals from the NCANDA dataset (60). Baseline resting-state fMRI data were pre-processed using the publicly-available NCANDA pipeline (158), which consisted of motion correction, outlier-detection, detrending, physiological noise removal as well as temporal (low pass frequency: 0.1, high pass frequency: 0.01) and spatial smoothing. Frames in individual rsfMRI time series were labeled as outliers if framewise displacement >0.3 mm/TR. After removing scans with usable frames *<* 7.8 min, each of the rs-fMRI images of the remaining 715 subjects (aged 12-21 years old, 52% female) were registered to the SRI24 atlas (159) and parcellated into 90 regions (80 cortical and 10 subcortical). The BOLD time series was normalized by the mean gray matter BOLD signal (pre-spatial and temporal filtering). We subsequently removed all censored frames, plus one frame before and two frames after, as well as uncensored segments of fewer than five contiguous frames. Using probabilistic tractography, structural connectomes (SC) were computed and parcellated into the same 90 regions.

Following the NCT analysis in the main text, we performed *k*-means clustering on BOLD time series data. Using a group-average SC, we calculated global and regional transition energies. We removed subjects whose global TE was +/-3 scaled MAD. We excluded subjects who exceeded alcohol, tobacco, marijuana or other drug usage based on criteria from (60). Further, we limited our cohort to only subjects who aged 15.9 years or younger to better match our ABCD cohort and limit the range of substance exposure. We performed 500 iterations of a matching algorithm to pair each FH+ subject with a FH-subject of the same sex and identified the match with the smallest average difference in age and mean framewise displacement. Our final cohort consisted of N = 64 subjects. We ran an ANCOVA on global TE including sex, age, family history of SUD, race/ethnicity, in-scanner motion, SES, MRI scanner model (GE vs Siemens), and the interactions terms sex:age, sex:family history of SUD, and family history of SUD:SES.

As in our main results, we find FH+ females > FH-females, and FH+ males < FH - males in mean global TE (Figure S9. While family history of SUD had a very small effect size (F= 0.03, *p* = 0.863), the interaction of sex and family history of SUD trended towards significance (F = 3.15, *p* = 0.082). Family history density was weakly positively and negatively correlated with mean global TE in females and males, respectively, though the correlations were not significant. Across regional TE’s, the majority of regions exhibited increased mean regional TE in FH+ versus FH-females, whereas the majority of regions had decreased TE in FH+ versus FH-males (Figure S10. We also replicated our previous result in the right pars orbitalis (“Frontal_Inf_Orb_R”), which was significant (pre-correction) for the interaction between sex and family history of SUD and exhibited the largest increase in TE of all regions in FH+ versus FH-females unpaired *t* tests. The bilateral middle occipital gyrus and the right precuneus (FH+ > FH-females, FH+ < FH-males) were also significant pre-correction for the interaction of sex and family history of SUD.

**Figure S9.**
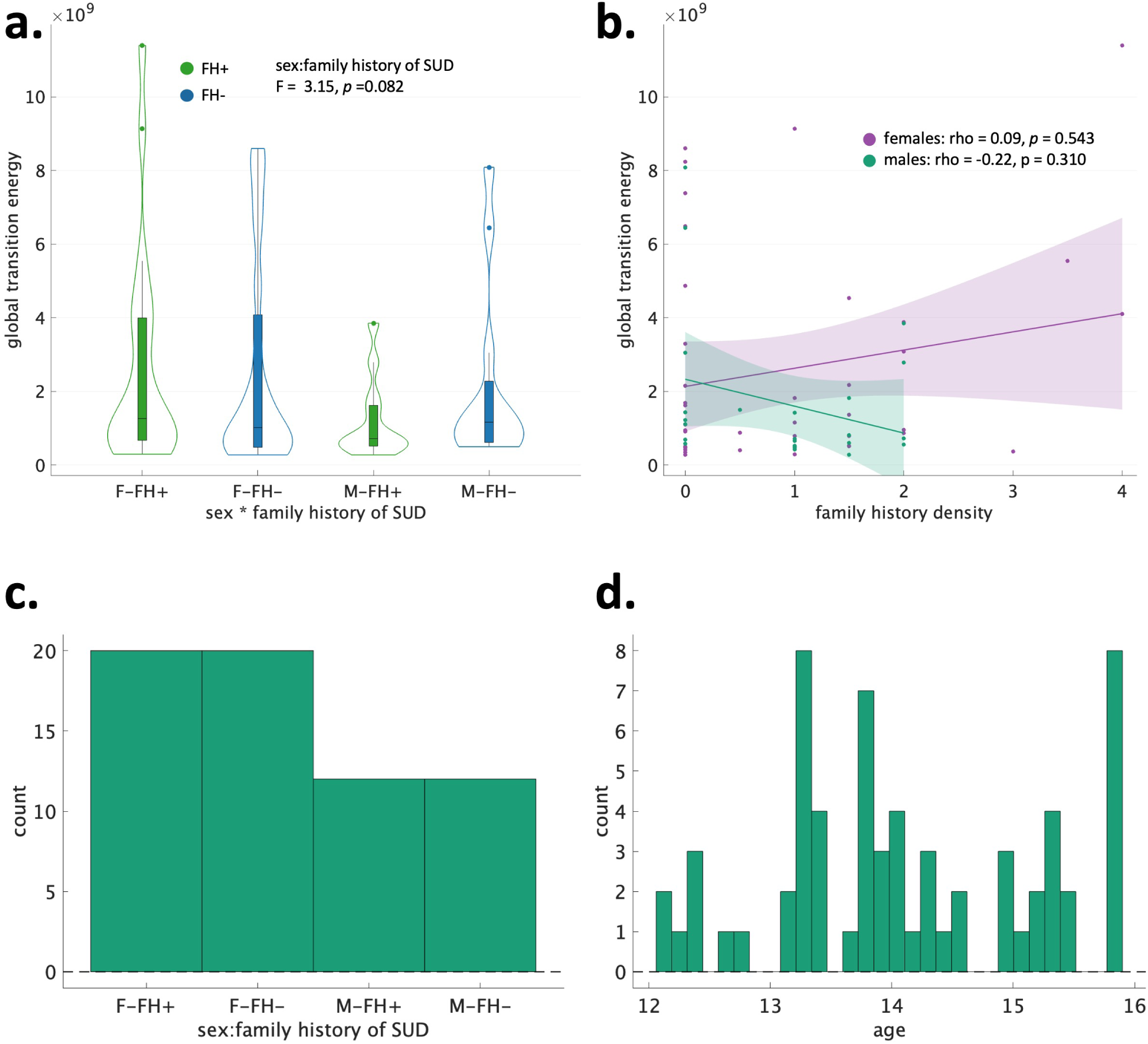
Replication of effect of interaction between sex and family history of SUD on global transition energy in external dataset (i.e., NCANDA).

**Figure S10.**
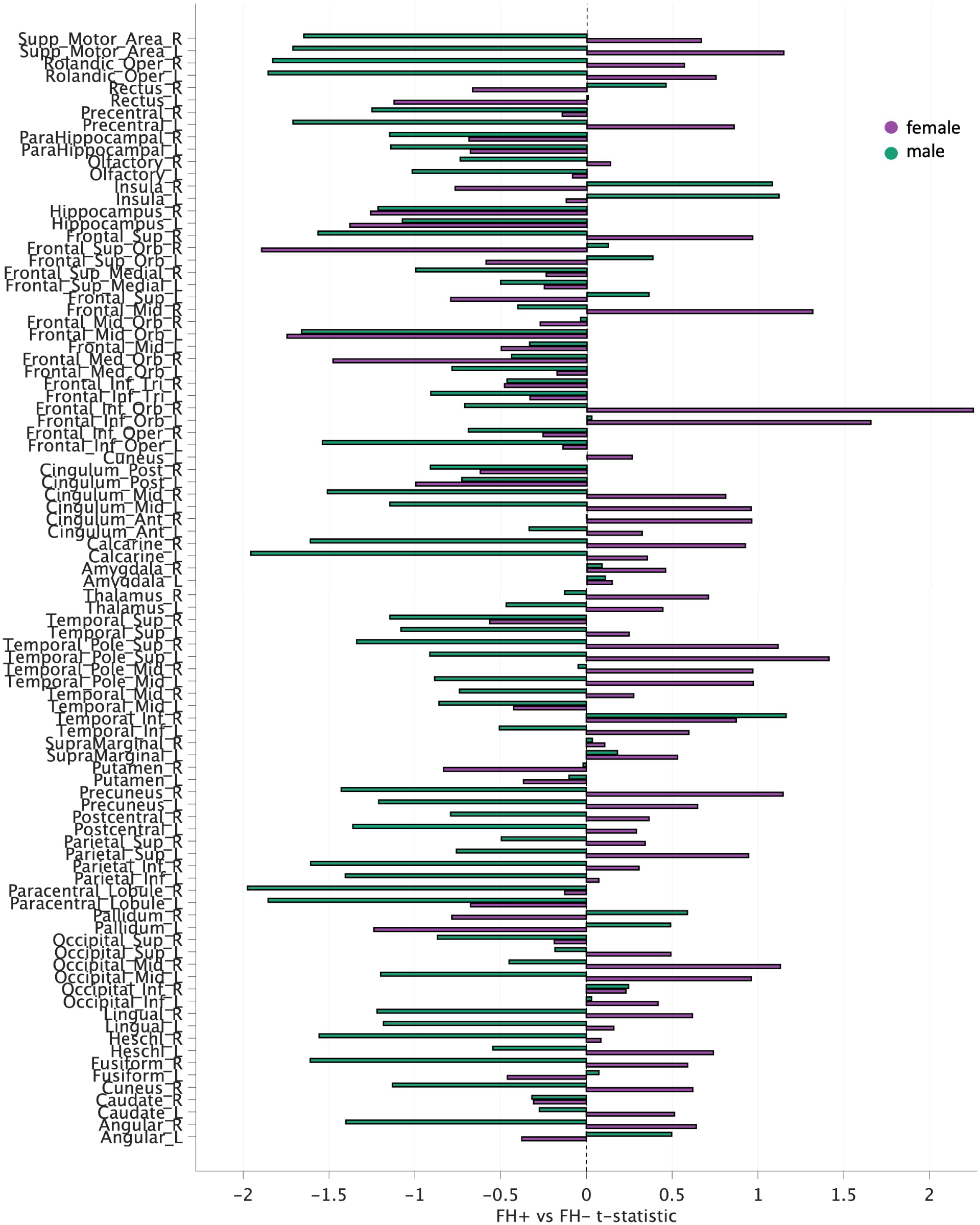
Replication of effect of interaction between sex and family history of SUD on regional transition energy in external dataset (i.e., NCANDA).

#### S0.5.3 Replication with individual structural connectomes in a cortex-only parcellation

We replicated our main results using individual strucutral connectomes (SC) rather than a group-average in order to determine whether our results are a result of individual differences in structural connecitivty. Individual structural connectomes (SC) were available only in the Desikan-Killiany atlas (68 cortical regions; DK68) for a subset of subjects from the ABCD study (N = 2080). See main text for details. After using the same exclusions described in the main text, 1710 subjects were included in analyses. *K*-means clustering (*k*=4) was run on these subjects’ regional resting-state functional MRI timeseries (DK68). Transition energies were calculated as described in the main text but with individual SCs (instead of a group-average SC) and only considering the seven Yeo Networks (59) in network-based analyses (not including subcortical or cerebellar networks). We find largely replicated results, except that family history-by-sex effects are significant in the VIS network is significant (pre-correction) and no longer significant in the VAT network. This effect in the VIS network was visible as a trend in group-average SC main analyses.

**Figure S11.**
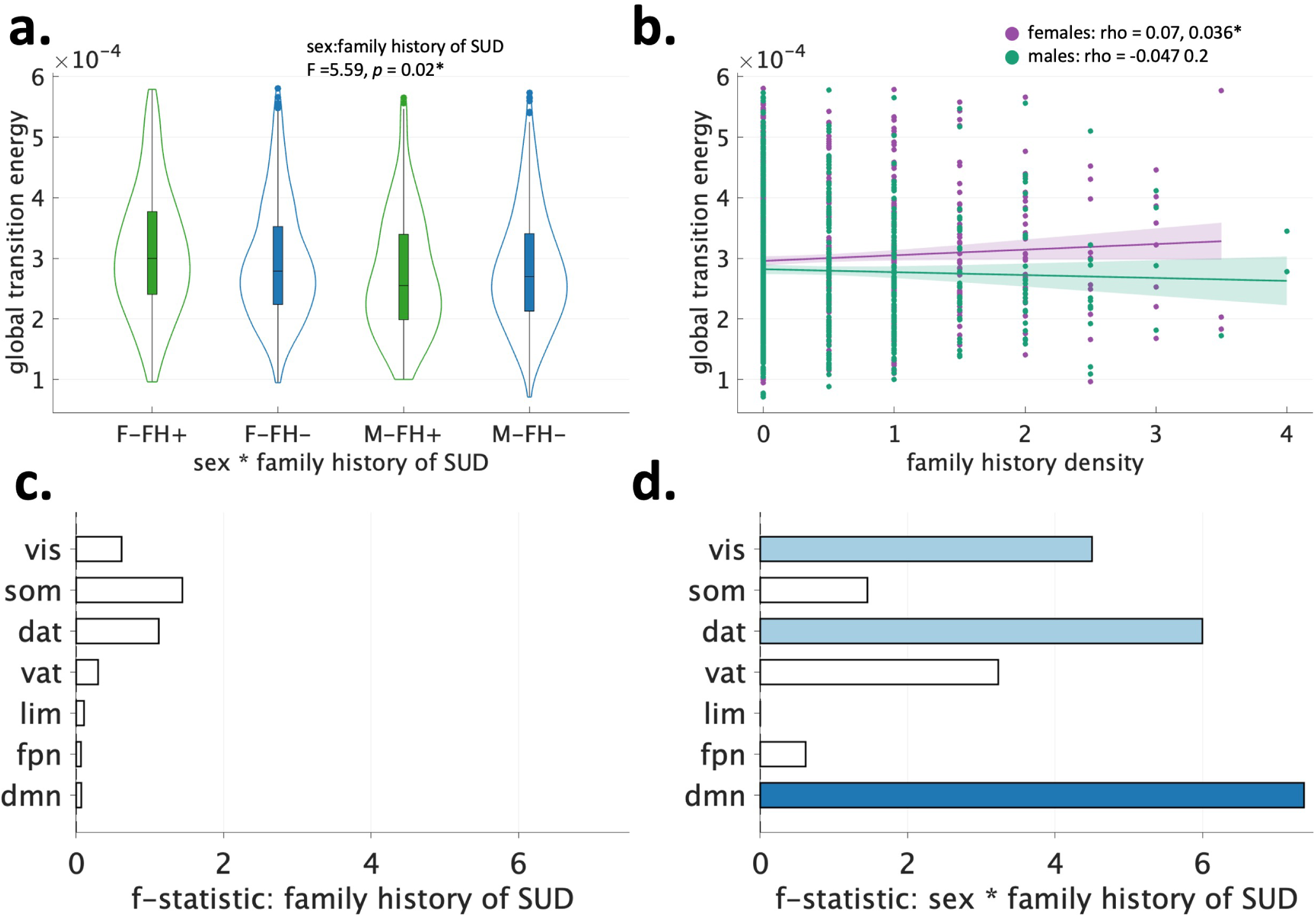
Global and network TE results are consistent with findings using group-average structural connectome, implicating functional dynamics in group differences.

### S0.6 Robustness analyses: stratified by single site, MRI models, and income levels

#### S0.6.1 Analysis within single-site

To demonstrate our findings are not the result of site effects, we analyzed subjects from within a single site. Site 16 was chosen due to its large sample size (N = 292) and previous work has noted site 16 has having “high-quality data” and was chosen as a reference site in a site-harmonization study (160). Site 16 utilized a Siemens Prisma MRI scanner. Using TE calculations from the analysis in the main text of *k*-means clustering (*k* = 4) from the entire cohort, we ran ANCOVAs on global and network TE values from subjects from site 16 only. Despite a relatively low N for FH+ subjects of either sex, we replicated the trends in global TE such that FH+ > FH- and FH+ < FH- and found a significant effect of the interaction of sex and family history. In network TE, we found these group-sex effects in the DMN and CER networks only.

**Figure S12.**
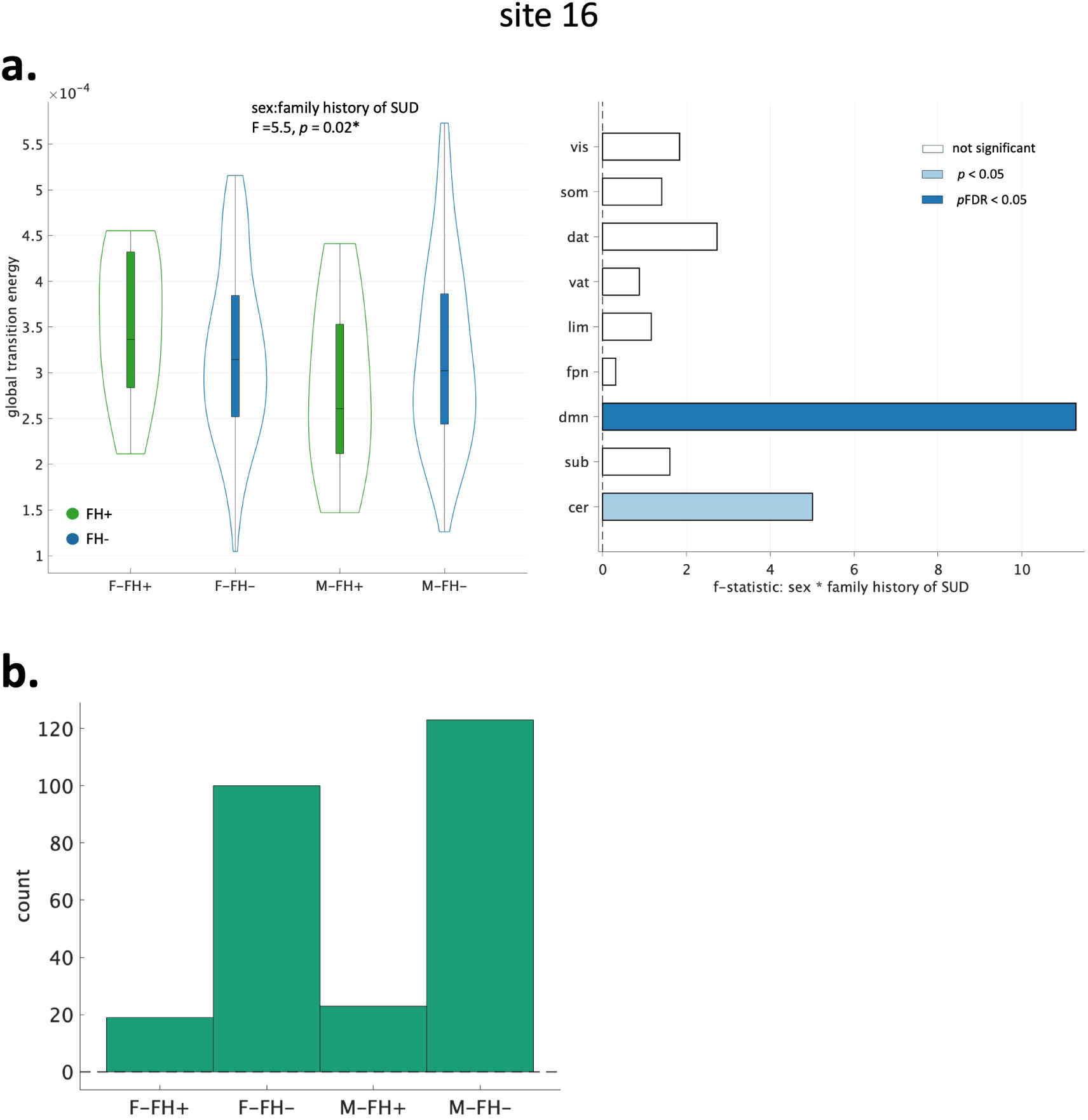

#### S0.6.2 Analyses within-MRI scanner models

Given the high effect size of MRI model on our NCT metrics, we ran ANCOVAs separately on subjects scanned on each of the three scanner MRI models utilized in our cohort: Siemens Prisma, Siemens Prisma Fit, and GE Discovery MR750. We find that our main results from the main text are driven by subjects scanned on Prisma scanner models (Siemens). In both Siemens models (Prisma and Prisma fit), we find results in global TE consistent with our main results, such that FH+ > FH- in females and FH+ < FH- in males. Within GE scanner subjects, global TE was also FH+ > FH-females, but differed in males (FH+ > FH-). However, only Prisma scanners showed a significant family history of SUD-by-sex for global TE. At the network level, our main results are partially replicated in Prisma and Prisma Fit scanners. Prisma scanner subjects exhibit significant (post-correction) effects of family history of SUD-by-sex in DMN and DAT networks, and pre-correction significance in VIS, LIM, FPN and SUB networks. Subjects from Prisma Fit scanners exhibit pre-correction significance in the VAT network. No networks are exhibited significant family history of SUD-by-sex effects in subjects from GE scanners.

As discussed in the main text, GE scanners had the smallest number of subjects, made up of younger subjects with higher levels of framewise displacement, lower household income, and more racially/ethnically diverse compared to the other two scanner models. The lack of replication of the results from our main text in GE scanner subjects may thus reflect differences in demographics or age-related differences in the neurodevelopmental trajectory. See Table **??** for subject demographics by MRI model. Furthermore, previous ABCD analyses have GE scanners to have lower-quality data more confounds and lacking real-time motion monitoring ((61; 62; 63; 64) compared to Siemens.

**Figure S13.**
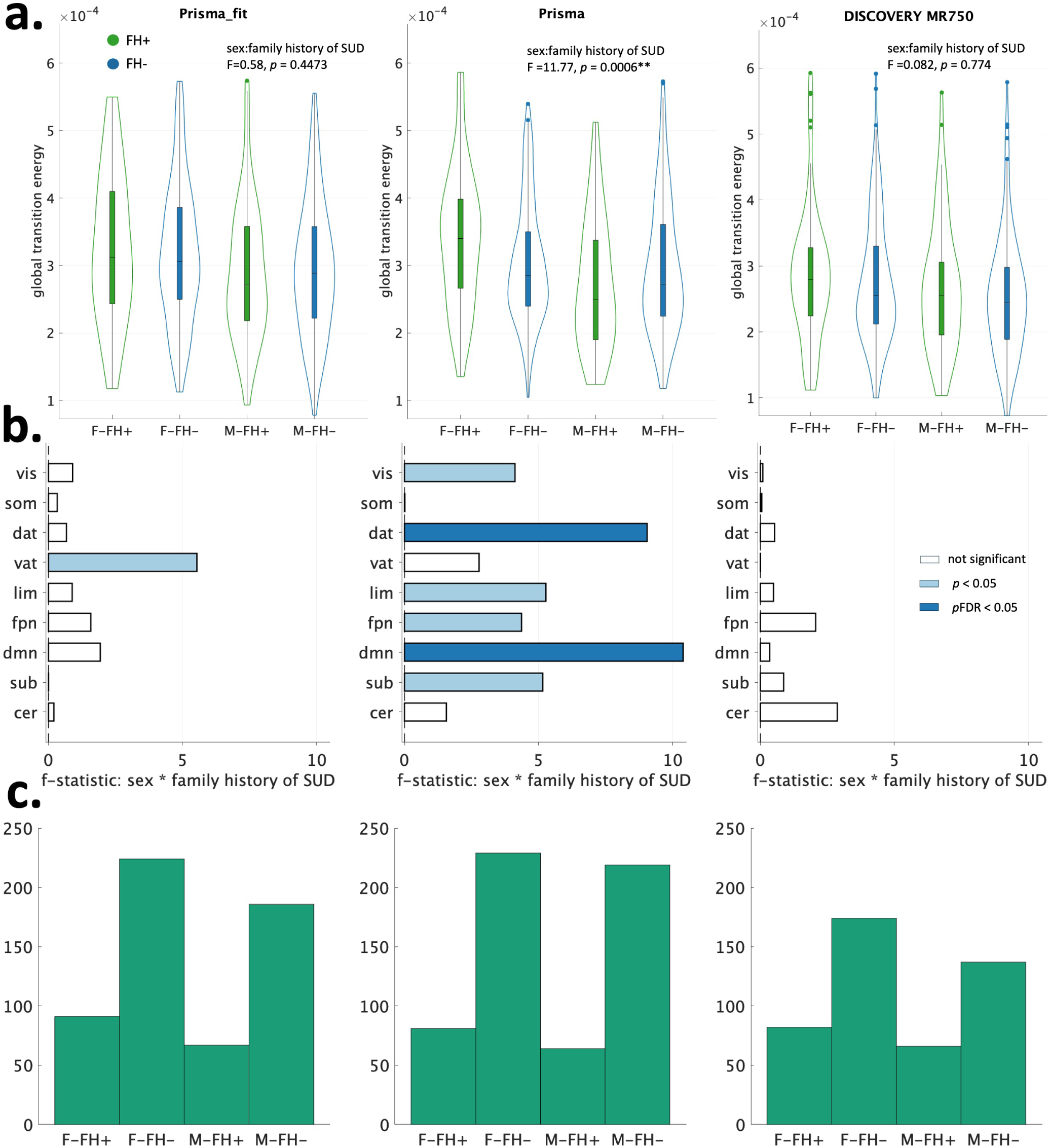

**Table S2.** Demographic and characteristic data of included subjects for the three MRI models (DISCOVERY MR750, Prisma_fit, and Prisma).

|  | DISCOVERY MR750 (n = 533) | Prisma_fit (n = 704) | Prisma (n = 710) |
| --- | --- | --- | --- |
| **FH Status** |  |  |  |
| FH+ | 148 (27.77%) | 161 (22.87%) | 147 (20.70%) |
| FH- | 314 (58.91%) | 429 (60.94%) | 462 (65.07%) |
| FH+/- | 71 (13.32%) | 114 (16.19%) | 101 (14.23%) |
| **Biological Sex, N(%)** |  |  |  |
| Male | 239 (44.84%) | 322 (45.74%) | 339 (47.75%) |
| Female | 294 (55.16%) | 382 (54.26%) | 371 (52.25%) |
| **Age in months, Mean (SD)** |  |  |  |
| Male | 119.33 (7.73) | 120.64 (7.53) | 121.02 (7.33) |
| Female | 118.76 (7.40) | 120.46 (7.19) | 120.29 (7.70) |
| **Framewise Displacement, Mean (SD)** |  |  |  |
| Male | 0.133 (0.091) | 0.130 (0.092) | 0.115 (0.071) |
| Female | 0.120 (0.077) | 0.111 (0.062) | 0.111 (0.067) |
| **Household Income, N(%)** |  |  |  |
| < 50,000 | 151 (28.33%) | 109 (15.48%) | 170 (23.94%) |
| 50,000–100,000 | 137 (25.70%) | 205 (29.12%) | 246 (34.65%) |
| 100,000+ | 245 (45.97%) | 390 (55.40%) | 294 (41.41%) |
| **Parent Education, N(%)** |  |  |  |
| > High School | 4 (0.75%) | 1 (0.14%) | 4 (0.56%) |
| High School/GED | 6 (1.13%) | 2 (0.28%) | 0 (0%) |
| Some College | 523 (98.12%) | 701 (99.57%) | 706 (99.44%) |
| Associates/Bachelor | 0 (0%) | 0 (0%) | 0 (0%) |
| Post-graduate | 0 (0%) | 0 (0%) | 0 (0%) |
| **Race/Ethnicity, N(%)** |  |  |  |
| White | 292 (54.78%) | 465 (66.05%) | 476 (67.04%) |
| Black | 34 (6.38%) | 67 (9.52%) | 44 (6.20%) |
| Hispanic/Latinx | 117 (21.95%) | 94 (13.35%) | 135 (19.01%) |
| Asian | 23 (4.32%) | 14 (1.99%) | 2 (0.28%) |
| Other | 67 (12.57%) | 64 (9.09%) | 53 (7.46%) |
| **In Utero Substance, N(%)** |  |  |  |
| Yes | 20 (3.75%) | 24 (3.41%) | 31 (4.37%) |
| No | 513 (96.25%) | 680 (96.59%) | 679 (95.63%) |
| **Parent Mental Health, N(%)** |  |  |  |
| Yes | 283 (53.10%) | 360 (51.14%) | 381 (53.66%) |
| No | 250 (46.90%) | 344 (48.86%) | 329 (46.34%) |

#### S0.6.3 Analyses within-income levels

Given the significant interaction of family history of SUD and income category on global TE, we investigated whether our findings were consistent across all the three income levels: 1, 2, or 3 (low to high). After *k*-means clustering (*k* = 4) and calculating TE values across the entire cohort, we ran an ANCOVA on data from each income category and display the results here. From the same variables included in ANCOVAs as described in the main text, we removed household income, parental education and the interaction term of family history of SUD and household income. We find our main results are primarily driven by the largest group of subjects in the high-income category (income group 3), which replicate both global effects (FH+ > FH-females, FH+ < FH-males) and family history-by-sex effects in the DMN and DAT. Individuals in the income group 1 show the same effect in females (FH+ > FH-) global TE, but also an increase in males (FH+ > FH-). Individuals in income group 2 show decreased global TE in males (FH+ < FH-) but a slight decrease in females as well. Income levels 1 and 2 have do not show significant family history-by-sex effects in any network. Studies have shown that in the ABCD cohort, which is made up of a majority of those with higher income, individuals from higher income families have higher curiosity and availability to substances, and earlier substance use initiation (65; 66; 67).

**Figure S14.**
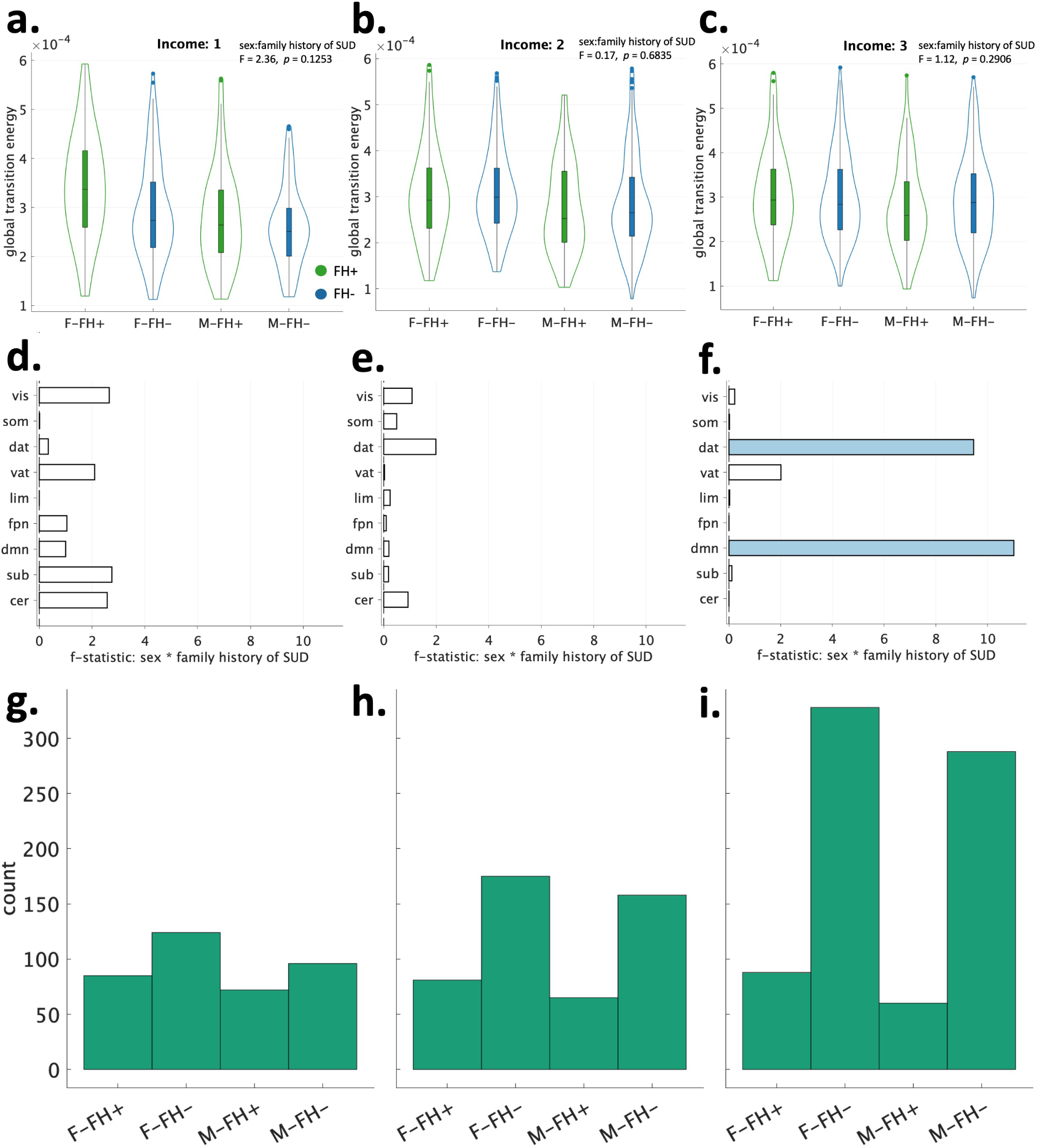

## Notes

### Competing Interest Statement

The authors have declared no competing interest.

